# Bitter taste receptors stimulate phagocytosis in human macrophages through calcium, nitric oxide, and cyclic-GMP signaling

**DOI:** 10.1101/776344

**Authors:** Indiwari Gopallawa, Jenna R. Freund, Robert J. Lee

## Abstract

Bitter taste receptors (T2Rs) are GPCRs involved in detection of bitter compounds by type 2 taste cells of the tongue, but are also expressed in other tissues throughout the body, including the airways, gastrointestinal tract, and brain. These T2Rs can be activated by several bacterial products and regulate innate immune responses in several cell types. Expression of T2Rs has been demonstrated in immune cells like neutrophils; however, the molecular details of their signaling are unknown. We examined mechanisms of T2R signaling in primary human monocyte-derived unprimed (M0) macrophages (MΦs) using live cell imaging techniques. Known bitter compounds and bacterial T2R agonists activated low-level calcium signals through a pertussis toxin (PTX)-sensitive, phospholipase C-dependent, and inositol trisphosphate receptor-dependent calcium release pathway. These calcium signals activated low-level nitric oxide (NO) production via endothelial and neuronal NO synthase (NOS) isoforms. NO production increased cellular cGMP and enhanced acute phagocytosis ∼3-fold over 30-60 min via protein kinase G. In parallel with calcium elevation, T2R activation lowered cAMP, also through a PTX-sensitive pathway. The cAMP decrease also contributed to enhanced phagocytosis. Moreover, a co-culture model with airway epithelial cells demonstrated that NO produced by epithelial cells can also acutely enhance MΦ phagocytosis. Together, these data define MΦ T2R signal transduction and support an immune recognition role for T2Rs in MΦ cell physiology.

## Introduction

Taste family 2 receptors (T2Rs) are GPCRs involved in bitter taste perception on the tongue [1, 2]. However, T2Rs are expressed throughout the body, including the nose and sinuses [3, 4] and lung [5]. T2Rs canonically decrease cAMP via Gα-gustducin (Gα_gust_) or Gα_i_ [6, 7] and Gβγ activation of phospholipase C (PLC), production of IP_3_, and elevation of calcium (Fig 1B) [3]. There are 25 T2Rs on the human tongue [1]. Nasal ciliated epithelial cells express T2Rs 4, 14, 16, 38, and possibly others. In addition to their canonical role in taste, mounting evidence suggests T2Rs are *bona fide* immune recognition receptors for chemosensation of bacteria. Cilia T2Rs detect bacterial products and activate local defense responses within seconds [7, 8], including Ca^2+^-dependent nitric oxide (NO) production that increases ciliary beating and has direct antibacterial effects [8, 9]. T2R38, T2R10, and T2R14 respond to bacterial acyl-homoserine lactone (AHL) quorum-sensing molecules [9-11]. T2Rs 4, 16, and 38 respond to *Pseudomonas* quinolone signal (PQS), and T2R14 responds to heptyl-hydroxyquinolone (HHQ) secreted by *Pseudomonas* [7].

**Fig. 1.**
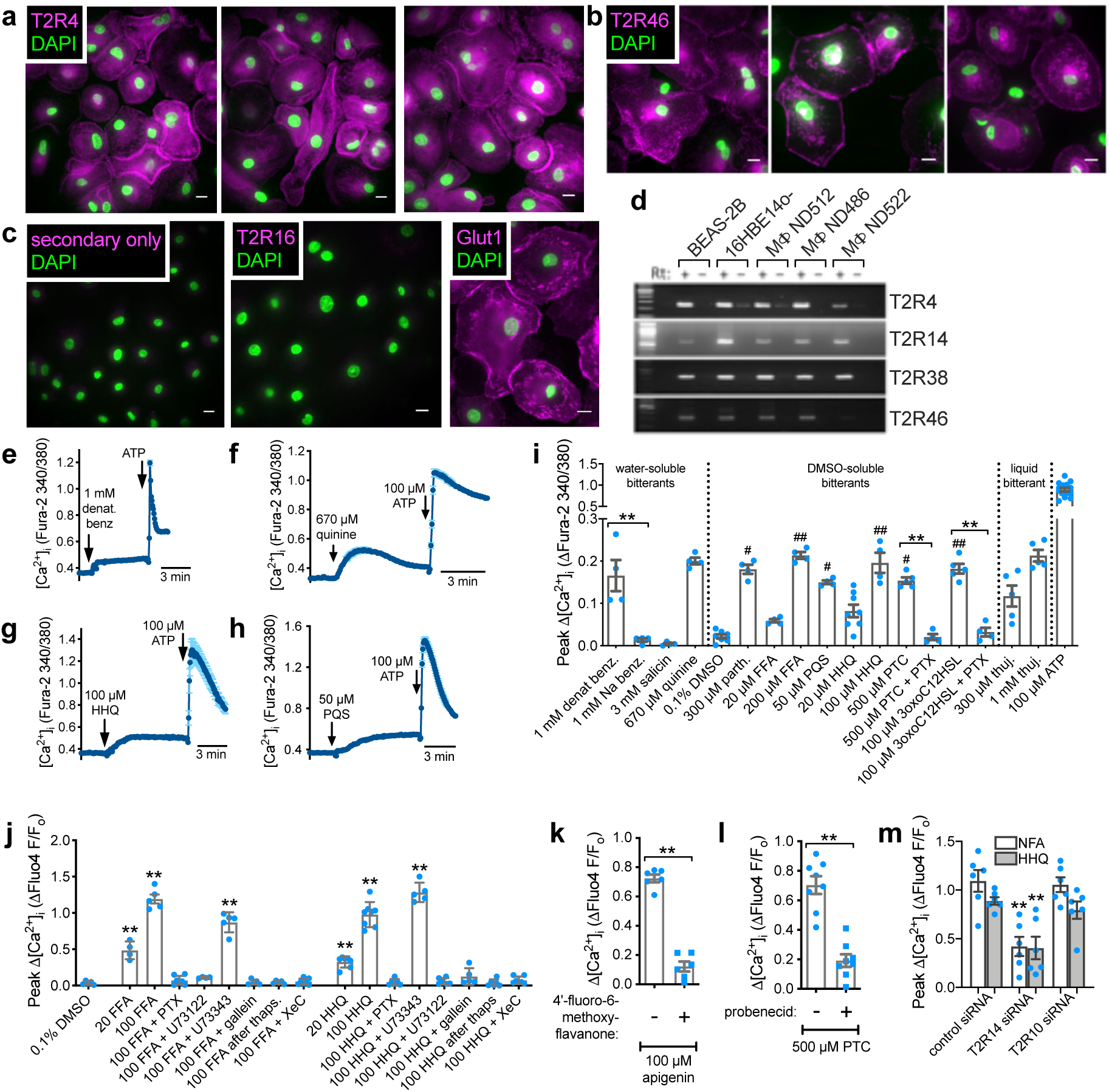
Unprimed (M0) monocyte-derived MΦs express functional bitter taste receptors (T2Rs). **a-c** Plasma membrane staining observed for T2R4 (*a*) and T2R46 (*b*) reminiscent of plasma membrane-localized glucose transporter GLUT1 (*c*). No immunofluorescence was observed with secondary only control or T2R16 antibody (*c*). **d** T2Rs detected in MΦs by rtPCR. Airway lines were used as control. **e-h** Representative low-level Ca^2+^ responses (fura-2) observed in response to denatonium benzoate (*e*), quinine (*f*), HHQ (*g*), and PQS (*h*). Subsequent stimulation with purinergic receptor agonist ATP used as control for viability. **i** Peak change in Ca^2+^ (fura-2 340/30 ratio) during 5 min stimulation with water soluble, DMSO-soluble, and liquid bitter compounds. ATP response shown for comparison. Also shown is inhibition of responses to PTC and 3oxoC12HSL by pertussis toxin (PTX; 100 ng/ml, 18 hrs pretreatment). Significance by one-way ANOVA with Bonferroni posttest with preselected paired comparisons; \*\**p* <0.01 between bracketed bars and ^#^*p* <0.05 and ^##^*p* <0.01 denote significance compared with DMSO only for DMSO-soluble bitterants. **j** Peak Ca^2+^ responses (fluo-4 F/F_o_) with T2R14 agonists FFA and HHQ (concentrations are µM) in the presence of inhibitors of GPCR signaling. Significance by one-way ANOVA, Dunnett’s posttest comparing each value to DMSO only. **k** Peak Ca^2+^ responses (fluo-4) to T2R14/39 agonist apigenin ± T2R14/39 antagonist 4’-fluoro-6-methoxy-flavanone (50 µM). Significance by Student’s t test; \*\**p* <0.01. **l** Bar graph showing peak Ca^2+^ (fluo-4) in response to T2R38 agonist PTC ± antagonist probenecid (1 mM). Significance by Student’s t test; \*\**p* <0.01. **m** Peak Ca^2+^ (fluo-4) during stimulation with T2R14 agonists NFA or HHQ in MΦs pre-treated with ON-TARGET plus SMARTpool siRNAs. Significance by one-way ANOVA with Bonferroni posttest; \*\**p* <0.01 vs control siRNA condition for each respective agonist. Representative traces are averages of 10-30 MΦs from single experiments. Data points in bar graphs are independent experiments using cells from ≥6 experiments (≥3 separate donors; ≥2 experiments per each donor)

Clinical importance of extraoral T2Rs in immunity is supported by correlation of polymorphisms rendering T2R38 non-functional with susceptibility to chronic rhinosinusitis CRS [9, 12-17]. A high number of T2R polymorphisms exist, contributing to individual taste preferences, but possibly also contributing to susceptibility to infection. It is critical to elucidate mechanisms of extraoral chemosensation by T2Rs to understand if and how to target T2Rs to activate innate immune responses.

Dedicated immune cells also express T2Rs [18-21]. T2R38 detects AHLs in neutrophils [18, 19] and is expressed in both resting and activated lymphocytes [20]. Circulating human monocytes, natural killer (NK) cells, B cells, T cells, and polymorphonuclear (PMN) leukocytes also express T2Rs [21]. Stimulation of PMNs with saccharin, which activates both T2Rs and T1R2/3 sweet receptors, enhanced leukocyte migration [21]. However, our understanding of the details of T2R signaling and physiological consequences in immune cells are limited.

Our goal was to determine if T2Rs in primary human macrophages (MΦs) are coupled to calcium and if this activates NO as in airway epithelial cells. MΦs are important players in early innate immune responses, and unprimed (M0) MΦs express Ca^2+^-activated endothelial (e) and/or neuronal (n) nitric oxide synthase (NOS) isoforms, though studies of e/nNOS in MΦ function are limited. The majority of studies of NO in MΦs focuses on higher level NO production via upregulation of inducible iNOS in LPS ± IFNγ-activated (classically-activated or M1) MΦs and the relationship of the NO to bacterial killing or metabolism [22]. However, iNOS is not expressed or is expressed at very low levels in monocytes and unstimulated M0 MΦs [22], while eNOS and/or nNOS are expressed [23-28] and can be activated by calcium [23, 26, 28].

Although relatively few papers examine e/nNOS activation in MΦ function, evidence suggests a potential role for NO in phagocytosis. Stimulation of Fcγ receptor (FcγR, not a GPCR) in M0 human MΦs activates both eNOS and nNOS to enhances phagocytosis [23], and the NO produced upregulates nNOS expression [23]. IFNγ also up-regulates phagocytosis in an NO-dependent manner in mouse MΦs [29]. Thus, we hypothesized that T2R activation of eNOS or nNOS may play a role in very early responses of MΦs encountering secreted “bitter” bacterial metabolites.

## Results

### MΦs express functional T2Rs tied to calcium signaling

Human monocytes obtained from healthy apheresis donors were differentiated to MΦs by adherence culture for 12 days, confirmed by functional expression of H1 receptors (**Supplementary Fig. 1**). We observed expression of several T2Rs in unprimed (M0) MΦs. Immunofluorescence using previously validated antibodies [7, 8] directed against T2R4 (**Fig. 1a**) and T2R46 (**Fig. 1b**), but not T2R16 (**Fig. 1c**), exhibited plasma membrane staining similar to GLUT1, a major MΦ glucose transporter [30]. Reverse transcription (rt) PCR for T2Rs 4, 14, 38, and 46 demonstrated expression in MΦs from 3 different individuals (**Fig. 1d**).

We tested a variety of T2R agonists (listed in **Supplementary Table 1**) using live cell calcium imaging. We observed low-level but sustained calcium responses to denatonium benzoate, which activates eight T2Rs including T2R4 and T2R46 (**Fig. 1e**) [31, 32], and to quinine, which activates 11 T2Rs including T2R14 and 46 (**Fig. 1f**) [31, 32]. Bacterial T2R agonists PQS and HHQ also increased calcium (**Fig. 1g-h**). A panel of bitterants revealed responses to T2R14 agonists thujone and flufenamic acid (FFA), T2R38 agonist phenylthiocarbamide (PTC), T2R14/38 agonist 3-oxo-dodecanoylhomoserine lactone (3oxoC12HSL), and T2R14/46 agonist parthenolide [31, 32]. Peak responses are summarized in **Fig. 1i** and representative traces in **Supplementary Fig. 2**. No response was observed with T2R16 agonist salicin, fitting with observed lack of expression (**Fig. 1c**). Stimulation of Gα_q_-coupled purinergic receptors with ATP was a positive control.

The responses to PTC were blocked by pertussis toxin (PTX; **Fig. 1i**), which ADP ribosylates and inactivates both Gα_gust_ and Gα_i_. Calcium responses to FFA and HHQ were inhibited by PTX and PLC inhibitor U73122, which inhibits T2R responses in airway cells [7, 8], Gβγ inhibitor gallein, which inhibits T2R responses in airway smooth muscle [33], and inositol trisphosphate receptor (IP_3_R) inhibitor xestospongin C (**Fig. 1j**). Moreover, responses to thujone, HHQ, or FFA were eliminated after intracellular calcium store depletion (thapsigargin in calcium-free solution) (**Fig. 1j**). Thus, T2R agonists activate calcium release from intracellular stores through GPCR signaling. Representative traces from experiments in **Fig. 1i-j** are in **Supplementary Fig. 2** and **3**. We found no differences in responses in MΦs differentiated by adherence + cytokine M-CSF (**Supplementary Fig. 4**).

To more concretely tie these results specifically to T2Rs, we used known T2R inhibitors; 4’-fluoro-6-methoxyflavanone is a T2R14 and T2R39 antagonist [34], while probenecid inhibits T2R16 and T2R38 [35]. Calcium responses to apigenin, which activates T2R14 and T2R39, were inhibited after pretreatment with 4’-fluoro-6-methoxyflavanone (**Fig. 1k**). Calcium responses to PTC were inhibited after pretreatment with probenecid (**Fig. 1l**). Representative experiments are in **Supplementary Fig. 5.** Knockdown of T2R14 with pooled siRNAs inhibited responses to T2R14 agonists niflumic acid (NFA; [31, 32]; **Fig. 1m**) and HHQ (**Fig. 1m**).

### MΦ T2R activation reduces baseline and stimulated cAMP signaling

As described above, T2R-stimulated Gα_i_ or Gα_gust_ can reduce cAMP through inhibition of adenylyl cyclase or activation of phosphodiesterase, respectively (**Fig. 2a**). Therefore, we also tested if bitter agonist stimulation reduced cAMP in MΦs. We utilized an mNeonGreen-based cAMP biosensor (cADDis [36]) in a baculovirus modified for mammalian cells (BacMam) to monitor cAMP in real time (**Fig. 2b**). Stimulation with β-adrenergic agonist isoproterenol increased cAMP (**Fig.2c**), whereas stimulation with 3oxoC12HSL, FFA, or PQS reduced cAMP (**Fig. 2d**). The cAMP reductions were eliminated by PTX (**Fig. 2d**). Moreover, cAMP increases with isoproterenol were reduced in the presence of PQS or 3oxoC12HSL (**Fig. 2e)**; this was likewise eliminated by PTX (**Fig. 2f**). We confirmed these results using another cAMP biosensor, the mTurquoise2-Venus FRET-based EPAC-S^H187^ [37]. MΦs expressing EPAC-S^H187^ exhibited PTX-sensitive cAMP decreases during stimulation with quinine, FFA, or 3oxoC12HSL (**Fig. 2g**).

**Fig. 2.**
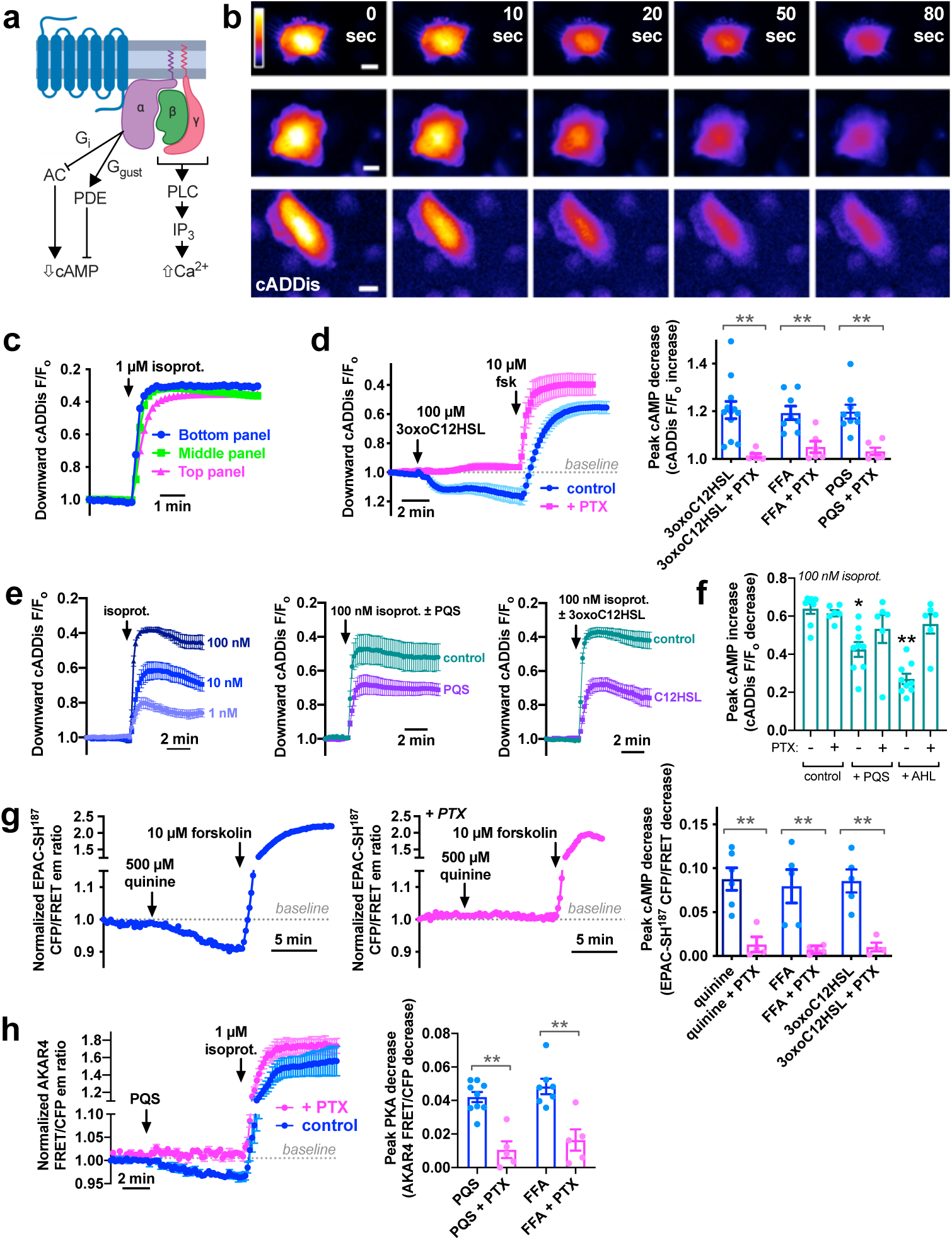
T2R signaling in MΦs decreases cAMP. **a** Diagram (created using Biorender.com) showing canonical T2R transduction pathway by which Gαi (G_i_) or Gα gustducin (G_gust_) decreases cAMP. **b-c** Representative image series from 3 separate experiments showing MΦs expressing green downward cADDis (b) and graphs (c) showing reproducibility of fluorescence changes in response to cAMP-elevating isoproterenol. Decrease in fluorescence equals an increase in cAMP, thus shown as an upward deflection (note inverse y axis). **d** Representative traces (left) and bar graph (right) showing cAMP decreases with 3oxoC12HSL, FFA, and PQS in control MΦs (blue) but not PTX-treated MΦs (pink); adenylyl cyclase-activating forskolin used as control. Traces are mean ± SEM of ≥6 independent experiments. **e** Traces of cADDis fluorescence (mean ± SEM; ≥6 independent experiments) during stimulation with isoproterenol ± PQS or 3oxoC12HSL. **f** Bar graph of peak cAMP increases with 100 nM isoproterenol ± PQS or 3oxoC12HSL (AHL) ± PTX. Graph shows mean ± SEM; each data point equals one independent experiment (n = ≥6 total from ≥3 donors). Significance by one-way ANOVA with Dunnett’s posttest (control is isoproterenol only, no PTX). **g** Traces (left and middle) of changes EPAC-S^H187^ CFP/FRET fluorescence emission ratio with quinine ± PTX. Each representative trace is a single experiment. Downward deflection equals decrease in cAMP. Right is bar graph of peak cAMP decreases with quinine, FFA, and 3oxoC12HSL ± PTX (mean ± SEM; ≥6 experiments each condition from ≥3 donors). **h** Trace of AKAR4 FRET/CFP fluorescence emission ratio (left; mean ± SEM of independent experiments)with PQS (100 µM) ± PTX. Downward deflection equals a decrease in PKA activity. Bar graph (right) shows peak PKA decrease with PQS or FFA ± PTX. Data points in bar graphs use cells from ≥3 donors (≥ 6 experiments total, ≥2 per donor). Significance in *d, g*, and *h* by one-way ANOVA with Bonferroni posttest. Significance in *f* by one-way ANOVA with Dunnett’s posttest comparing values to control (no PTX); \**p*<0.05 and \*\**p*<0.01

Data demonstrate that T2Rs affect both basal and stimulated cAMP levels in MΦs. Using a ratiometric Cerulean-Venus FRET-based PKA biosensor, AKAR4 [38], we observed that PKA activity was also reduced during stimulation with PQS or FFA, and this was eliminated by PTX (**Fig. 2h**). Together, these data indicate that both arms of canonical T2R signaling appear to be active in MΦs when stimulated with bitter compounds.

### MΦ T2R calcium responses drive NO production

These T2R-activated low-level calcium responses were reminiscent of airway cells [3, 7, 8]. We tested if NO production was also downstream. M0 MΦs express calcium-sensitive eNOS and/or nNOS [23-28]. We confirmed eNOS transcript by rtPCR (**Fig. 3a**) and detected eNOS and nNOS by Western (**Fig. 3b**). We used the fluorescent indicator DAF-FM to monitor production of reactive nitrogen species (RNS) in living MΦs in real time. DAF-FM terminally reacts with NO and RNS derivatives, increasing fluorescence. Non-specific NO donor S-nitroso-N-acetyl-D,L-penicillamine (SNAP) was used as a positive control.

**Fig. 3.**
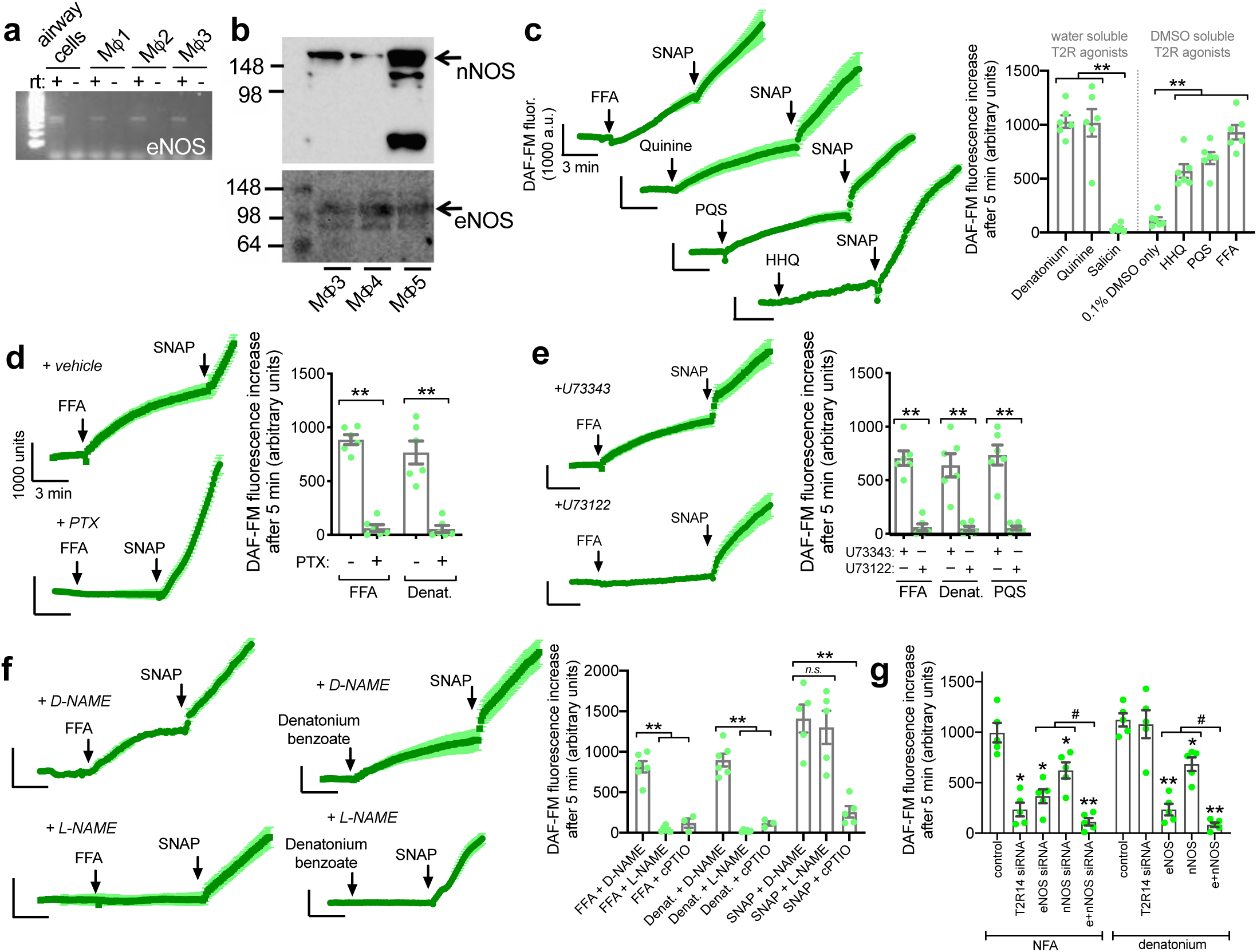
T2R stimulation activates NO production in MΦs. **a** Reverse transcription (rt) PCR of MΦs from 3 donors showing expression of eNOS (NOS3). **b** Westerns of eNOS and nNOS using MΦ lysates from 3 donors. **c** Traces and bar graph showing DAF-FM fluorescence increases during stimulation with FFA (100 µM), quinine (500 µM), PQS (100 µM), HHQ (100 mM), or salicin (3 mM). Non-specific NO donor SNAP (10 µM) shown as a control. **d** Traces and bar graph showing DAF-FM fluorescence increase in response to FFA (100 µM) or denatonium benzoate (1 mM) ± PTX. **e** Traces and bar graph showing DAF-FM fluorescence increases in response to FFA (100 µM), denatonium benzoate (1 mM), or PQS (100 µM) in the presence of PLC inhibitor U73122 or inactive analog U73343 (30 min pretreatment, 10 µM). **f** Traces and bar graph showing DAF-FM fluorescence increases in response to FFA (100 µM) or denatonium benzoate (1 mM) in the presence of L-NAME or inactive D-NAME (100 µM; 30 min pretreatment) as well as with NO scavenger cPTIO (10 µM). **g** DAF-FM increases in response to 100 µM NFA or 1 mM denatonium benzoate in MΦs treated with Accell SMARTpool siRNAs as indicated. Traces are mean ± SEM from 20-30 MΦs from single representative experiments. Bar graphs are mean ± SEM with data points shown from independent experiments using cells from ≥3 donors (≥6 experiments total, ≥2 per donor). Significance determined by one-way ANOVA with Bonferroni posttest; **p>0.01 vs control, ^#^p<0.05 vs bracketed group, *n.s.* = no statistical significance

Several T2R agonists, but not T2R16 agonist salicin, caused increases in DAF-FM fluorescence (**Fig. 3b**) that were blocked by PTX (**Fig. 3c**) or U73122 (**Fig. 3d**), suggesting they are downstream of T2R calcium signaling. DAF-FM fluorescence increases in response to T2R stimulation, but not SNAP, were blocked by pretreatment with NOS inhibitor L-N^G^-niroarginine methyl ester (L-NAME), but not inactive D-NAME, and were reduced in the presence of NO scavenger carboxy-PTIO (cPTIO; **Fig. 3e**). DAF-FM results were confirmed by measuring NO decomposition products NO_2_^-^ and NO_3_^-^ in culture media (**Supplementary Fig. 6**). Thus, DAF-FM fluorescence increases reflect NOS activation.

### MΦ T2R signaling increases cGMP

To test the hypothesis that NO production would increase cGMP, we utilized a fluorescent cGMP biosensor (Green GENIe). We observed increases in cGMP production with denatonium benzoate or parthenolide (**Fig. 4a**), which activate calcium and NO responses, but not with salicin or sodium benzoate, which do not activate calcium or NO. The cGMP responses to FFA or PQS were blocked by L-NAME or cPTIO (**Fig. 4b**), PTX (**Fig. 4c**), or elimination of calcium (**Fig. 4d**). The cGMP increase was enhanced when MΦs were co-infected with BacMam to over express soluble guanylyl cyclase (sGC), further tying these responses to NO-induced cGMP.

**Fig. 4.**
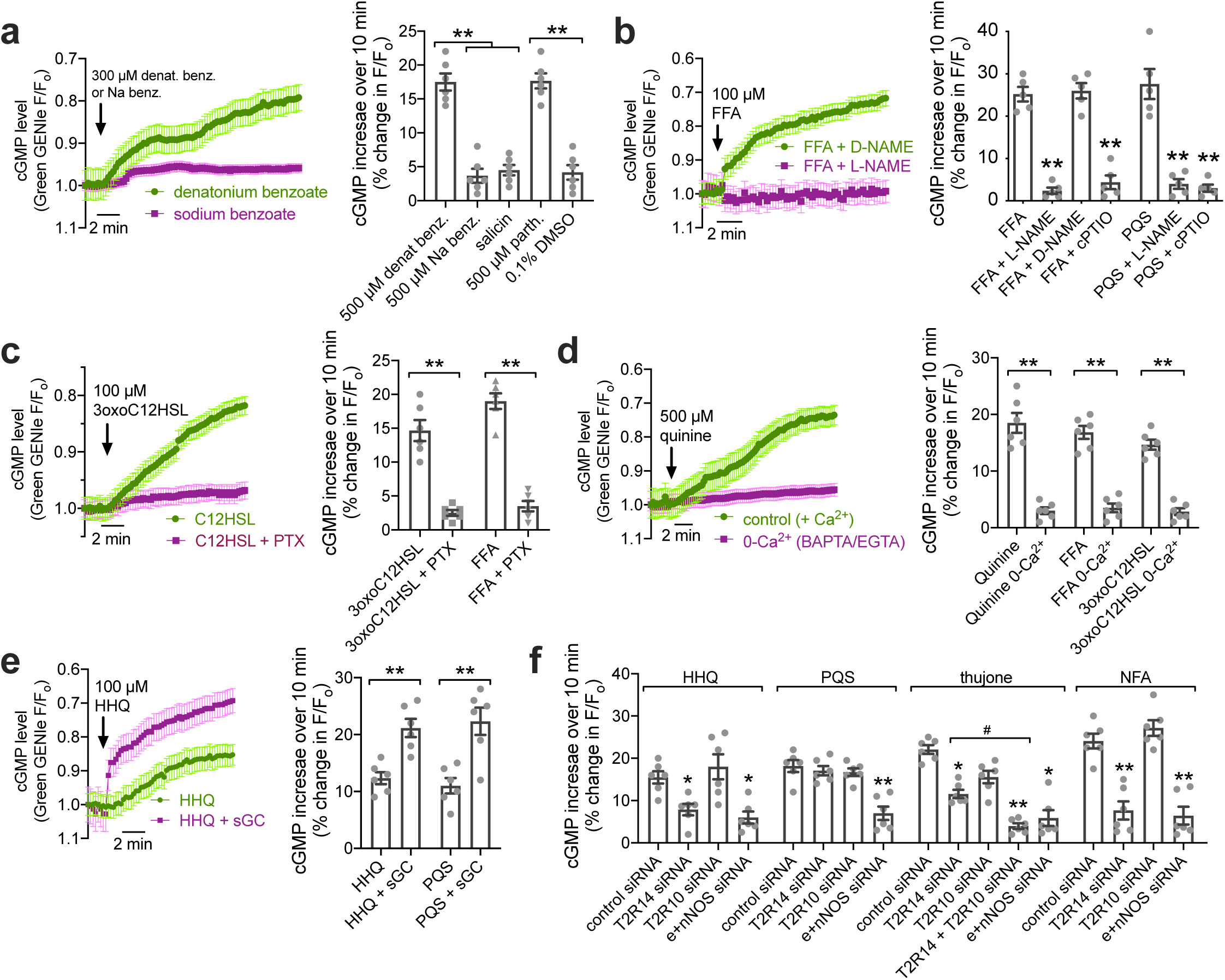
T2R-induced NO production increases cGMP. **a** Traces and bar graph showing changes in Green GENIe cGMP indicator with T2R agonists. Decrease in fluorescence equals increase in cGMP, plotted as upward deflection (note inverse y axis). **b** Traces and bar graph of cGMP increases with FFA or PQS (100 µM each) ± L-NAME or D-NAME (100 µM 30 min pretreatment) or cPTIO (10 µM). **c** Traces and bar graph of cGMP increases with 3oxoC12HSL or FFA (100 µm each) ± PTX (100 ng/ml pretreatment for 18 hrs). **d** Traces and bar graph of cGMP increases with quinine, FFA (100 µM), or 3oxoC12HSL (100 µM) ± Ca^2+^ signaling; 0-Ca^2+^ conditions are 1 µM BAPTA-AM loading for 60 min with stimulation in HBSS containing no added Ca^2+^ and 1 mM EGTA. **e** Traces and bar graph of cGMP increases with HHQ or PQS (100 µM each) ± co-infection with BacMam expressing soluble guanylyl cyclase (sGC). **f** Bar graph of cGMP increases in response to HHQ (100 µM), PQS (100 µM), thujone (600 µM), or NFA (100 µM) in MΦs pre-incubated with ON-TARGET plus SMARTpool siRNAs directed against T2R14 or T2R10, a cocktail of eNOS and nNOS, or control siRNA. All traces are mean ± SEM of ≥6 independent experiments. Bar graphs are mean ± SEM with data points shown from independent experiments using cells from at least 2 separate donors (≥ 5 experiments total, ≥2 experiments per donor). Significance determined by one-way ANOVA with Bonferroni posttest; * or ^#^*p*<0.05 and \*\**p*<0.01

We used siRNA to confirm involvement of T2Rs. MΦs treated with control siRNA exhibited cGMP responses to HHQ, PQS, thujone, and niflumic acid (NFA; **Fig. 4f**). Treatment with a cocktail of eNOS and nNOS siRNA blocked the response to each agonist. T2R14 siRNA, but not T2R10 siRNA significantly reduced the cGMP responses to T2R14 agonists HHQ and NFA, but the response to PQS, which does not activate T2R14 [7], remained intact (**Fig. 4f**). The response to thujone, which activates both T2R14 and T2R10 [31, 32], was slightly inhibited by either T2R14 or T2R10 siRNAs, and more fully inhibited with a cocktail of both siRNAs (**Fig. 4f**).

### MΦ T2R activation may alter MΦ metabolism through combined elevations of calcium and NO

We briefly examined if T2R stimulation had acute effects on MΦ metabolic state, tied to MΦ activation [39], by live-cell imaging of NAD(P)H autofluorescence [40]. Ultraviolet autofluorescence increased with bitter stimulation in a PTX- and L-NAME-sensitive manner (**Supplementary Fig. 7**), suggesting an increase in NAD(P)H levels. Increases in autofluorescence were not observed with either thapsigargin or SNAP separately to non-specifically increase Ca^2+^ or NO, respectively. However, addition of thapsigargin plus SNAP robustly increased NAD(P)H autofluorescence, suggesting this is an effect of raising both Ca^2+^ and NO simultaneously (**Supplementary Fig. 7**). Together, data suggest that T2R stimulation may acutely alter MΦ metabolism, at least short term, supporting a role for T2Rs in activation of MΦ immune responses.

### MΦ T2R-driven NO signaling enhances phagocytosis

Activation of T2Rs in MΦs increases intracellular calcium to activate e/nNOS and produce NO and cGMP. What consequences does this have for MΦ cell physiology? As mentioned above, NO was implicated in regulation of phagocytosis. We tested if T2R stimulation increased acute phagocytosis of FITC-labeled *E. coli* bioparticles. Incubation of MΦs with FITC-*E. coli* in the presence of T2R bitter agonists that activated calcium and NO also increased phagocytosis by ∼300% (**Fig. 5a-c)**. Responses to FFA were inhibited by L-NAME, cPTIO, PKG inhibitor KT5823, or PTX (**Figure 5C**). Responses to denatonium benzoate were inhibited by PLC inhibitor U73122 or Gβγ inhibitor gallein (**Fig. 5d**). A cocktail of pooled T2R14 siRNA inhibited responses to NFA or HHQ but not denatonium benzoate, which does not activate T2R14 (**Fig. 5e**). Responses to all three agonists were reduced with e/nNOS siRNAs (**Fig. 5e**).

**Fig. 5.**
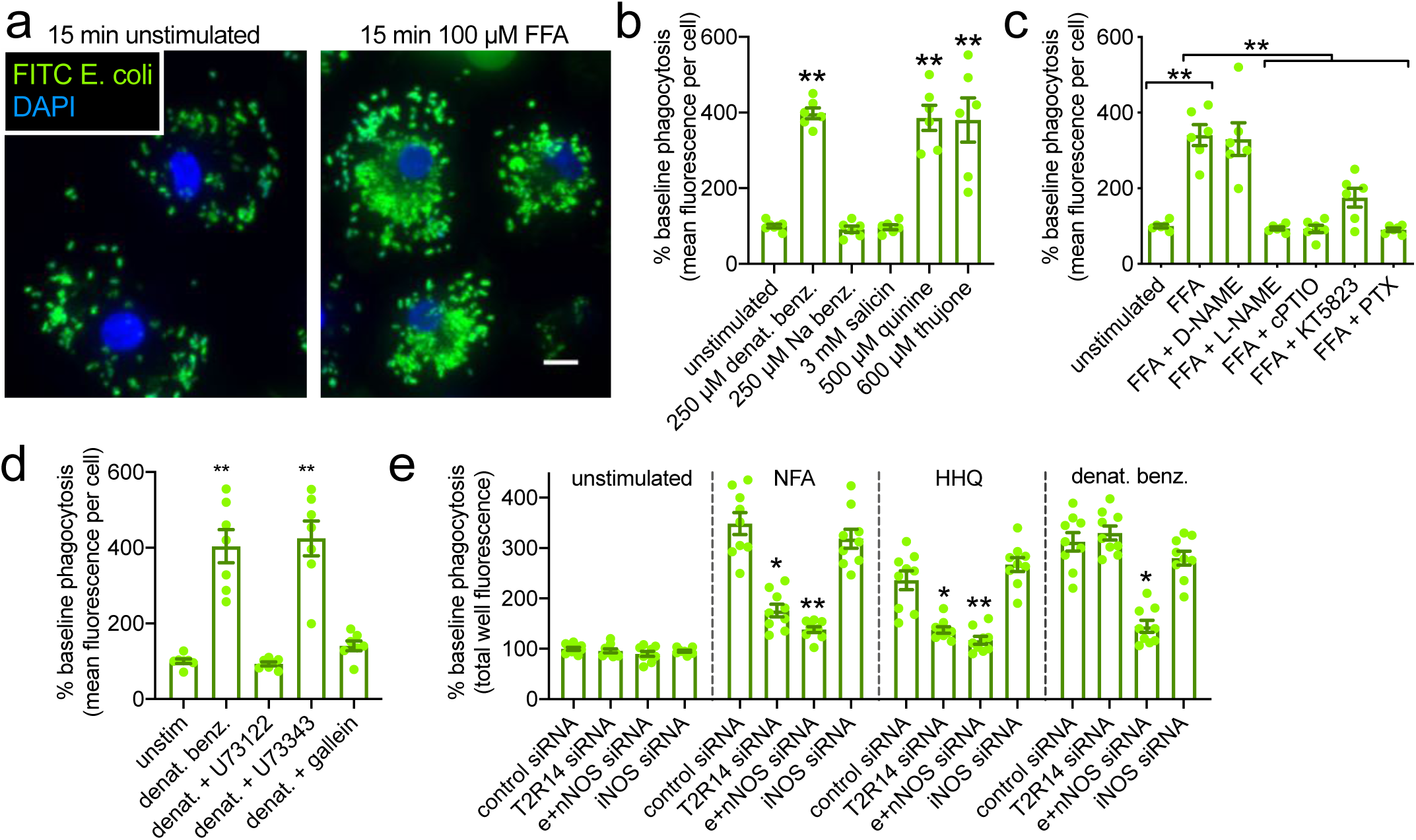
T2R-induced NO/cGMP acutely increases MΦ phagocytosis of FITC-labeled *E. coli*. **a** Representative image of fixed MΦs after 15 min with FITC *E. coli* ± FFA. **b** Bar graphs of normalized phagocytosis (quantified by microscopy) ± various T2R agonists. **c** Normalized phagocytosis (quantified by microscopy) ± FFA (100 µM) ± D/L-NAME (100 µM, 30 min pretreatment), cPTIO (10 µM), KT5823 (1 µM), or PTX (100 ng/ml; 18 hrs pretreatment). **d** Bar graphs showing normalized phagocytosis ± denatonium benzoate (1 mM) ± U73122 (10 µM, 30 min pretreatment), U73343 (10 µM, 30 min pretreatment) or gallein (100 µM). **e** Bar graph of phagocytosis ± NFA (100 µM), HHQ, (100 µM), or denatonium benzoate (1 mM) in MΦs treated with Accell SMARTpool siRNAs as indicated. Results in *b, c*, and *d* were quantified by microscopy; each independent experiment is average of 10 fields from a single well. Results in *e* were quantified by plate reader; each independent experiment is average of 2 wells. Bar graphs are mean ± SEM with data points shown from independent experiments using cells from ≥3 separate donors (≥2 independent experiments per donor)

We confirmed that we were observing phagocytosis using *Staphylococcus aureus* bioparticles labeled with pHrodo, a dye that fluoresces at the low pHs existing in lysosomes and phagosomes (**Fig. 6a**). MΦs phagocytosed pHrodo *S. aureus*, evidenced by increased fluorescence, when incubated at 37 °C but not 4 °C; this was enhanced by denatonium benzoate but not sodium benzoate (**Fig. 6b and d**). Similar results were observed with FFA (**Fig. 6c and e**). Increased phagocytosis with FFA was inhibited by L-NAME, cPTIO (**Fig. 6c and e**), U73122, or PTX (**Fig. 6f**). Increased phagocytosis with denatonium benzoate or quinine was also inhibited by PKG inhibitor KT5823 but not PKA inhibitor H89 (**Fig. 6g**).

**Fig. 6.**
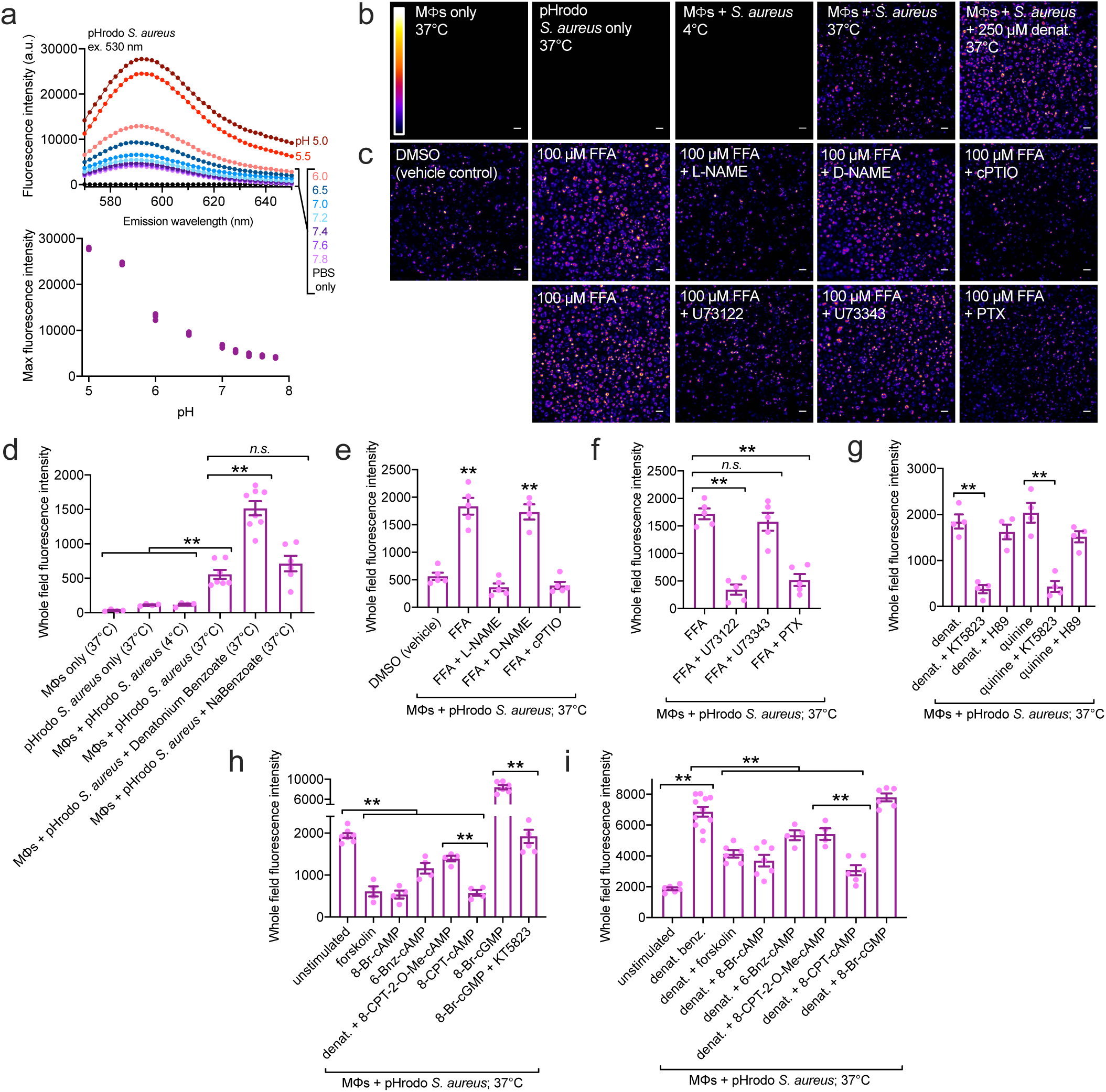
T2R-induced NO and cGMP acutely increases MΦ phagocytosis of pHrodo-labeled *S. aureus*. **a** pHrodo-labeled S aureus were resuspended in PBS buffered to various pHs as indicated to confirm increase in fluorescence with decreasing pH. Experiment in top graph representative of 3 independent replicates, plotted on bottom graph. **b** Representative images (20x; scale bar 20 µm) of pHrodo labeled *S. aureus* and MΦs showing phagocytosis occurring only when both were combined and only at 37°C. **c** Representative images of increased phagocytosis with FFA and inhibition with L-NAME (100 µM, 30 min pretreatment), cPTIO (10 µM), U73122 (10 µM, 30 min pretreatment), and PTX (100 ng/ml; 14 hrs pretreatment). **d** Quantification of experiments performed as in *b*. Significance by Bonferroni posttest; ** *p*<0.01. **e-f** Quantification of experiments performed as in *c*. **g** Similar experiments were performed with denatonium benzoate (2 mM) and quinine (500 µM); phagocytosis increase was inhibited by KT5823 but not H89 (5 µM, 10 min preincubation each). Significance by one-way ANOVA with Dunnett’s posttest; ** *p*<0.01. **h** Quantification of pHrodo *S. aureus* phagocytosis assays ± 10 µM forskolin or cell permeant cAMP or cGMP analogs as indicated. Forskolin or cAMP analogs inhibited phagocytosis; however 8-Br-cGMP increased phagocytosis via PKG, as it was blocked by KT5823 (5 µM, 10 min preincubation). **i** Quantification of pHrodo S. aureus assays with 1 mM denatonium benzoate ± forskolin or cAMP analogs or cGMP analog. Significance by 1-way ANOVA with Bonferroni posttest. Data points in *d*-*i* are independent experiments (≥6 from ≥2 individual donors, all taken with identical microscope settings), each experiment is average of 10 fields from a single well. Incubations for *b*-*g* were 20 min. Incubations for *h* and *i* were 60 min

Others have suggested that cAMP inhibits MΦ phagocytosis or activation [41-44]. We hypothesized that T2R-mediated cAMP decreases also facilitate phagocytosis. Baseline phagocytosis was inhibited by direct adenylyl cyclase activation by forskolin or cell permeant cAMP analogs, including 8-Br-cAMP and 8-CPT-cAMP, which activate both PKA and EPAC, and to a lesser extent by PKA-specific 6-Bnz-cAMP or EPAC-specific 8-CPT-2-O-Me-cAMP (**Fig. 6h**). In contrast, cell permeant cGMP analog 8-Br-cGMP increased phagocytosis via PKG; this was inhibited by KT5823. Thus, increased cAMP likely reduces phagocytosis partially through PKA and partially through EPAC, unlike cGMP which increases phagocytosis through PKG. Denatonium benzoate-induced phagocytosis was inhibited by forskolin or permeant cAMP analogs used above, but were not inhibited by 8-Br-cGMP (**Fig. 6i**). Together, these data suggest both the calcium/NO and cAMP arms of the T2R pathway facilitate MΦ phagocytosis.

### NO crosstalk from airway epithelial cells may also enhance MΦ phagocytosis

Airway epithelial cells also make NO in response to various stimuli including T2R [3, 7] and estrogen receptor activation [45]. We hypothesized that MΦs close to the airway surface may be influenced by local intercellular NO production. We designed an assay to test epithelial-MΦ crosstalk using PTX-treated MΦs and H441 small airway epithelial cells, which produce NO in response to 17β-estradiol (E2) (**Supplementary Fig. 8**). Although MΦs exhibited low-level calcium transients to 10 nM E2, these were eliminated by PTX pretreatment (**Supplementary Fig. 9**).

DAF-FM-loaded, PTX-treated MΦs were incubated with unloaded H441s separated by a transwell filter in close proximity (≤1 mm; **Fig. 7a**). Stimulation of H441 cells with E2 resulted in increased MΦ DAF-FM fluorescence (**Fig. 7b**), suggesting NO produced by the H441s translated to an increase in RNS within the MΦs. Thus, epithelial NO/RNS could act as an intercellular signal. Inclusion of cPTIO or pretreatment of H441 cells with PTX or L-NAME blocked the MΦ DAF-FM response (**Fig. 7c**), but when MΦs were pretreated with L-NAME, there was no inhibition (**Fig. 7c**). Thus, the NO/RNS originated in the H441 cells. E2 had no effect when MΦs were incubated in the absence of H441s (**Fig. 7c**). Together, these data support the potential of airway epithelial cells to produce enough NO to be sensed by MΦs in close proximity.

**Fig. 7.**
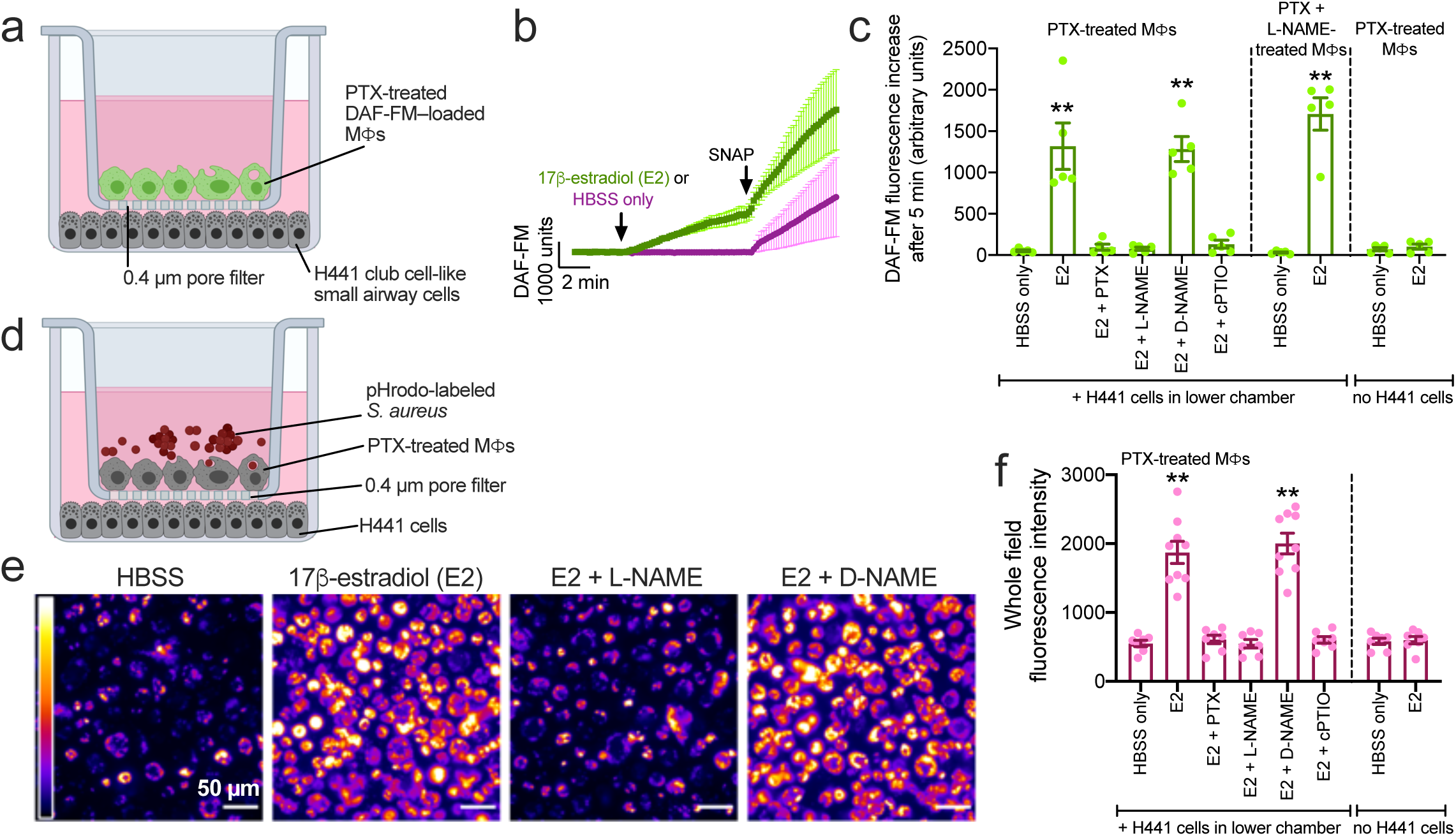
Airway epithelial cells can acutely increase MΦ phagocytosis via intercellular NO. **a** Diagram of experimental design for *b*-*c*. PTX-pretreated DAF-FM-loaded MΦs were placed in close proximity (<1 mm) to unloaded H441s separated by a permeable plastic filter. **b** Stimulation of H441s with 10 nM 17β-estradiol (E2) resulted in an increase in MΦ DAF-FM fluorescence. SNAP (10 µM) added at the end as a positive control. **c** Bar graph of DAF-FM increases from experiments performed as in *a*-*b*. DAF-FM increases were inhibited by treatment of H441s with PTX (100 ng/ml; 18 hrs) or L-NAME (10 µM, 30 min) or in the presence of cPTIO (10 µM). Treatment of MΦs with L-NAME did not alter responses, and there was no response to E2 in the absence of H441s. **d** Diagram of experimental design for *e*-*f*. PTX-pretreated MΦs incubated with pHrodo-labeled *S. aureus* were placed in close proximity (<1 mm) to H441s. **e** Representative images of fluorescence increases with PTX-pretreated MΦs with H441s stimulated as indicated for 30 min. **f** Bar graph of MΦs fluorescence increases quantified by microscopy with stimulations as indicated. E2 increased MΦ phagocytosis that was blocked by PTX, L-NAME, cPTIO, or absence of H441s. Bar graphs are mean ± SEM with data points shown from independent experiments using cells from at least 2 individual donors (≥2 independent experiments per donor). Significance by one-way ANOVA with Bonferonni posttest with each bar compared with its respective HBSS only control; ** p<0.01. Figures in *a* and *d* created with Biorender.com

To test if airway epithelial NO production could act as a signal to stimulate MΦ phagocytosis, we measured changes in pHrodo *S. aureus* phagocytosis by PTX-treated MΦs when H441s were stimulated with E2 (**Fig. 7d**). E2 stimulation of H441s increased MΦ phagocytosis (**Fig. 7e and f**), and this was blocked by H441 pretreatment with PTX or L-NAME or inclusion of cPTIO in the media (**Fig. 7f**). E2 had no effect when PTX-treated MΦs were incubated in the absence of H441 cells (**Fig. 7f**).

## Discussion

We only understand the “tip of the iceberg” of extraoral T2Rs and their signaling. We demonstrated that T2Rs in MΦs may enhance phagocytosis through calcium-driven NO-activation of guanylyl cyclase to increase cGMP (**Fig. 8**). Interestingly, this is a similar pathway to airway epithelial cells [3, 7-9, 46], but the physiological output is different. In ciliated cells, the pathway controls cilia beating; in MΦs, it regulates phagocytosis. Both processes are critical for innate defense, supporting a role for T2Rs as immune receptors. In MΦs encountering bitter bacterial agonists at sites of infection, activated T2Rs may “rev up” acute phagocytic activity while likely simultaneous toll-like receptor (TLR) stimulation up-regulates expression of iNOS and antimicrobial proteins such as lysozyme to further combat infection. Of note, T2Rs also respond to a variety of natural plant compounds associated with complementary medicines, including flavonoids [8, 47-49], as well as common clinical drugs [50, 51]. Activation of extraoral T2Rs in MΦs may underlie some effects of homeopathic plant-based treatments or off-target drug effects.

**Fig. 8.**
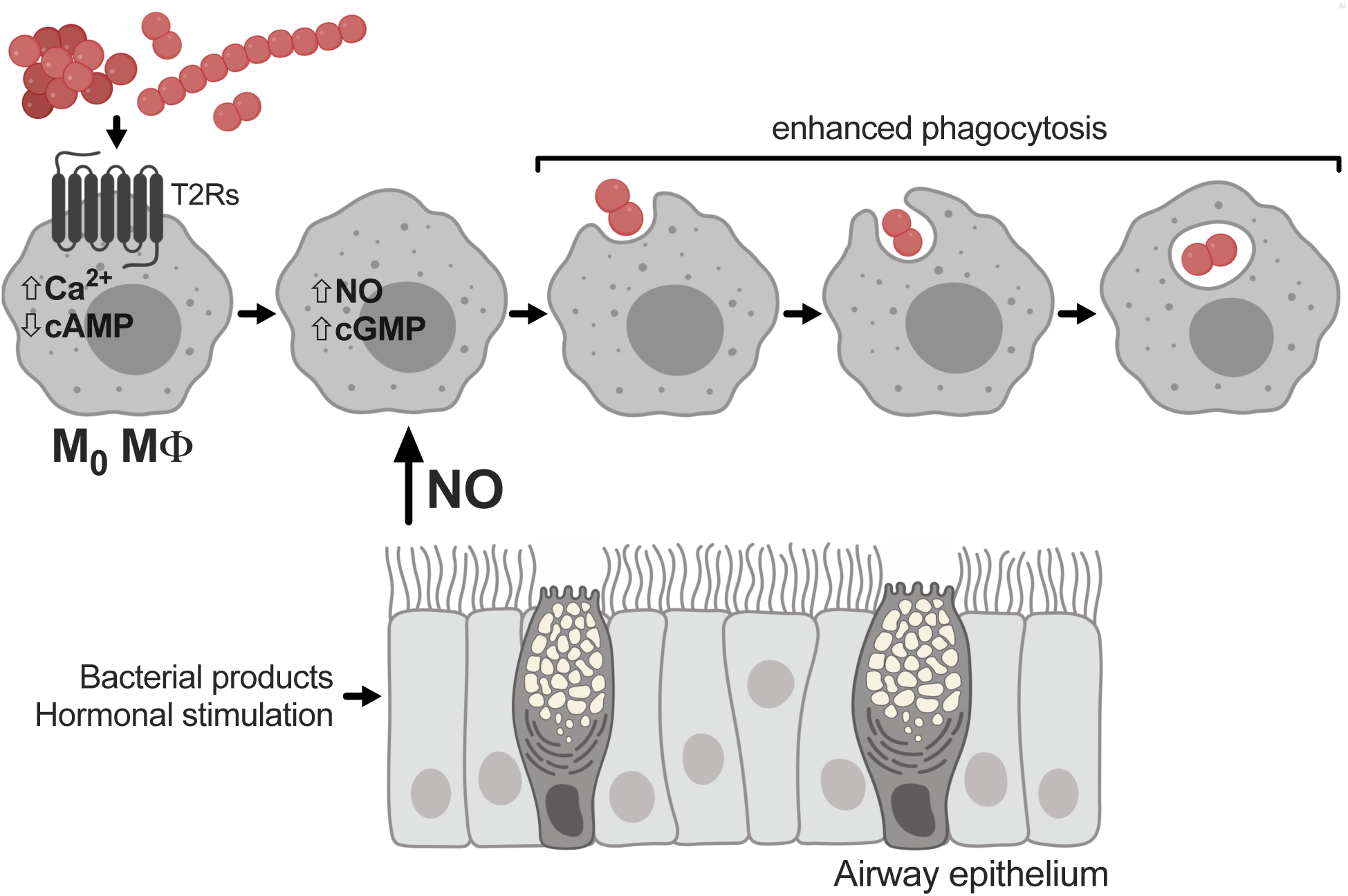
Model of T2R signaling in M0 MΦs. Activation of T2Rs increases Ca^2+^, lowers cAMP, activates eNOS and nNOS, increases cGMP, and enhances phagocytosis. Acute increase in phagocytosis may also be activated by airway epithelial cell NO production. Figure created with Biorender.com

This study also suggests that NO produced by airway epithelial cells could increase MΦ phagocytosis in a paracrine fashion (**Fig. 8**). NO produced during activation of T2Rs in airway epithelial cells [3, 7] may influence immune cells or amplify immune responses. NO may regulate phagocytosis via several mechanisms, including ADP-ribosylation of actin to regulate cytoskeletal polymerization, and pseudopodia formation, and phagocytosis in mouse peritoneal MΦs [52]. In Raw 264.7 mouse MΦ-like cells, NO acutely regulates actin organization through cGMP in combination with Ca^2+^/calmodulin [53], and cGMP/PKG may be important in phagocytic responses during TLR2 stimulation in THP-1 monocytic leukemia cells [54].

It is important to better understand e/nNOS and cGMP signaling in MΦs beyond phagocytosis. Production of cGMP was linked to protection against toxicity of peroxynitrite (ONOO-) or oxidized LDL [55], reduction of cytokine release during LPS stimulation [56] or high fat exposure [57, 58], and promotion of wound healing [59]. NO itself may be anti-inflammatory by activation of PPARγ, antagonizing NOX2 production of reactive oxygen species [60], or attenuation of NF-ΦB and activation of Nrf2 transcription factors [61]. However, MΦs from eNOS knockout mice exhibit reduced NO production, NF-ΦB activation, and iNOS upregulation during LPS stimulation [28].

Interestingly, T2R-induced lowering of cAMP levels may be another mechanism to enhance phagocytosis; cAMP elevation through prostaglandins decreases phagocytosis in human MΦs [41] and other inhibitory effects of cAMP have been reported [42-44]. Canonical T2R signaling is driven by Ca^2+^ downstream of Gβγ activation of PLC. The decrease in cAMP by G_gust_ was proposed to accentuate calcium signaling via type III IP_3_Rs in taste cells [62, 63], but data supporting this mechanism are limited. Increased cAMP and PKA phosphorylation of IP3R is more often reported to enhance calcium signaling [64], though PKA phosphorylation of type III IP3R has been suggested to slow kinetics of calcium release in pancreatic acinar cells [65]. If taste cell signaling is primarily driven by calcium [66], why have T2Rs not evolved to couple to G_q_ instead of G_gust_ or G_i_? This may be because, as observed here, decreases in cAMP have important biological effects in extraoral tissues.

## Materials and Methods

### Reagents and solutions

Unless indicated, all reagents and solutions were as described [7, 8, 67, 68]. Full details of materials, reagents, and solution compositions are in the **Supplementary Material**. Bitter agonists used and known cognate T2Rs are in **Supplementary Table 1**. Immunofluorescence (IF) microscopy was carried out as described [7, 8, 67, 68], with more details in the **Supplementary Material**.

### MΦ culture

Monocytes were isolated by the University of Pennsylvania Human Immunology Core from healthy apheresis donors with institutional review board approval by RosetteSep™ human monocyte enrichment cocktail (Stem Cell Technologies, Vancouver, Canada), and MΦs were differentiated by 12-days adherence culture in RPMI 1640 media containing 10% human serum and 1x cell culture pen/strep antibiotic mix (Gibco). All cells were deidentified before receipt. No investigator had contact with personal identifying information or demographics. Between 5-15 million cells were obtained from each individual. Cells from 29 individuals were used; some individuals donated multiple times during the course of the study. Details of siRNA protocols are in the **Supplementary Material**.

### Live cell imaging of intracellular calcium, reactive nitrogen species (RNS) production, and cAMP/cGMP signaling

Intracellular calcium and RNS were imaged using Fura-2 or Fluo-4 and DAF-FM, respectively, as described [7, 8, 67]. MΦs were infected with BacMams containing green downward cADDis or green downward GENie (both from Montana Molecular, Bozeman MT). For AKAR4 or Epac-S^H187^, MΦs on 8-well chambered coverglasses were transfected with 0.5 µg/well plasmid using Effectene (MilliporeSigma, Burlington, MA) per the manufacturer’s protocol. Full details are in the **Supplementary Material**.

### Phagocytosis assays

For microscopy, MΦs on chambered coverglasses were incubated with heat-killed FITC-labeled *E. coli* bioparticles at 250 µg/ml (strain K-12; Vybrant phagocytosis assay kit; ThermoFisher Scientific) in phenol red-free, low glucose DMEM ± bitter agonists or inhibitors for 15 min at 37°C. For microscopy, extracellular FITC was quenched with trypan blue and cells were washed ≥5x in PBS to remove residual extracellular FITC-*E. coli*. Remaining adherent MΦs were fixed in 4% formaldehyde (Electron Microscopy Sciences) for 10 min followed by DAPI staining. For pHrodo Red-labeled *S. aureus* (strain Wood 46; ThermoFisher A10010), living MΦs on chambered coverglass were visualized by fluorescence microscopy at room temperature (which drastically reduces any further phagocytosis) immediately after the assay incubation at 37°C.

For plate reader assays, MΦs on microplates were incubated similarly as above with 500 µg/ml FITC-*E. coli*. As phagocytosis was eliminated at 4 °C and almost completely eliminated at room temperature, we recorded fluorescence from living cells at room temperature immediately after the 15 min incubation with FITC-*E. coli*. Extracellular FITC was quenched with trypan blue, and fluorescence recorded on a Spark 10M plate reader (Tecan; 485 ex, 535 em). Phagocytosis assays were carried out similarly using 125 µg/ml pHrodo-*S. aureus* for 30 min at 37°C.

### Airway epithelial cell culture and coculture

H441 cells (ATCC HTB-174) were cultured in MEM with Earle’s salts (Gibco) containing 10% FetalPlex (Gemini Biosciences) in T75 flasks, lifted with trypsin when 75% confluent and transferred to 24 well plates and grown to confluence for MΦs co-culture experiments or seeded onto glass chamber slides and imaged for acute calcium and NO measurements.

For phagocytosis and NO co-culture, MΦs were seeded separately on transparent Falcon filters (#353095; 0.3 cm^2^; 0.4 µm pores), which come very close (0.8 mm) to the bottom of standard 24 well plates. Filters containing 100 µL HBSS on the apical side were transferred into the 24 well plates containing H441 cells and a small amount of HEPES-buffered HBSS (200 µL) at the time of the assay. MΦs were pretreated with PTX (100 ng/ml) for ∼18 hrs in media (37°C, 5% CO_2_), followed by copious washing with HBSS to remove residual PTX before co-incubation with H441s. Similarly, DAF-FM loading of MΦs was performed prior to co-incubation with H441s. Addition of fluorescently-labeled bacteria to MΦs on transwells was done at the time of assay start. Immediately after, 100 µL of a 3x E2 solution in HBSS was added to the basolateral side to stimulate H441 NO production and the 24 well plate was then incubated at 37°C.

### Data analysis and statistics

Statistical significance was determined in GraphPad Prism as indicated in figure legends; *p* <0.05 was considered statistically significant. Other analyses were performed in Excel. Data are presented as mean ± SEM. Data points in bar graphs are ≥3 independent experiments using cells from ≥2-3 donors.

## Acknowledgements

This study was supported by National Institutes of Health grants (R01DC016309, R21AI137484). Content is solely the responsibility of the authors and does not represent official views of the National Institutes of Health. The authors thank J. Riley (University of Pennsylvania Human Immunology Core, supported by P30-CA016520 and P30-AI045008) for access to monocytes and M. Victoria (University of Pennsylvania) for excellent assistance with MΦ culture and molecular biology and helpful comments on the manuscript. We thank N. Cohen (University of Pennsylvania) for sharing reagents. The authors have no conflicts of interest to declare.

## References

1 Behrens M and Meyerhof, W (2010) Oral and extraoral bitter taste receptors. Results Probl. Cell Differ. 52:87-99 doi: 10.1007/978-3-642-14426-4_8

2 Lee RJ and Cohen, NA (2015) Taste receptors in innate immunity. Cell. Mol. Life Sci. 72:217-236 doi: 10.1007/s00018-014-1736-7

3 Freund JR and Lee, RJ (2018) Taste receptors in the upper airway. World J Otorhinolaryngol Head Neck Surg 4:67-76 doi: 10.1016/j.wjorl.2018.02.004

4 Carey RM and Lee, RJ (2019) Taste receptors in upper airway innate immunity. Nutrients 11 doi: 10.3390/nu11092017

5 An SS and Liggett, SB (2017) Taste and smell gpcrs in the lung: Evidence for a previously unrecognized widespread chemosensory system. Cell. Signal. doi: 10.1016/j.cellsig.2017.02.002

6 Kim D, Woo, JA, Geffken, E, An, SS and Liggett, SB (2017) Coupling of airway smooth muscle bitter taste receptors to intracellular signaling and relaxation is via galphai1,2,3. Am. J. Respir. Cell Mol. Biol. 56:762-771 doi: 10.1165/rcmb.2016-0373OC

7 Freund JR, Mansfield, CJ, Doghramji, LJ, Adappa, ND, Palmer, JN, Kennedy, DW, Reed, DR, Jiang, P and Lee, RJ (2018) Activation of airway epithelial bitter taste receptors by pseudomonas aeruginosa quinolones modulates calcium, cyclic-amp, and nitric oxide signaling. J. Biol. Chem. 293:9824-9840 doi: 10.1074/jbc.RA117.001005

8 Hariri BM, McMahon, DB, Chen, B, Freund, JR, Mansfield, CJ, Doghramji, LJ, Adappa, ND, Palmer, JN, Kennedy, DW, Reed, DR, Jiang, P and Lee, RJ (2017) Flavones modulate respiratory epithelial innate immunity: Anti-inflammatory effects and activation of the t2r14 receptor. J. Biol. Chem. 292:8484-8497 doi: 10.1074/jbc.M116.771949

9 Lee RJ, Xiong, G, Kofonow, JM, Chen, B, Lysenko, A, Jiang, P, Abraham, V, Doghramji, L, Adappa, ND, Palmer, JN, Kennedy, DW, Beauchamp, GK, Doulias, P-T, Ischiropoulos, H, Kreindler, JL, Reed, DR and Cohen, NA (2012) T2r38 taste receptor polymorphisms underlie susceptibility to upper respiratory infection. J. Clin. Invest. 122:4145-4159

10 Lossow K, Hubner, S, Roudnitzky, N, Slack, JP, Pollastro, F, Behrens, M and Meyerhof, W (2016) Comprehensive analysis of mouse bitter taste receptors reveals different molecular receptive ranges for orthologous receptors in mice and humans. J. Biol. Chem. 291:15358-15377 doi: 10.1074/jbc.M116.718544

11 Jaggupilli A, Singh, N, Jesus, VC, Duan, K and Chelikani, P (2018) Characterization of the binding sites for bacterial acyl homoserine lactones (ahls) on human bitter taste receptors (t2rs). ACS Infect Dis 4:1146-1156 doi: 10.1021/acsinfecdis.8b00094

12 Adappa ND, Howland, TJ, Palmer, JN, Kennedy, DW, Doghramji, L, Lysenko, A, Reed, DR, Lee, RJ and Cohen, NA (2013) Genetics of the taste receptor t2r38 correlates with chronic rhinosinusitis necessitating surgical intervention. Int Forum Allergy Rhinol 3:184-187

13 Adappa ND, Zhang, Z, Palmer, JN, Kennedy, DW, Doghramji, L, Lysenko, A, Reed, DR, Scott, T, Zhao, NW, Owens, D, Lee, RJ and Cohen, NA (2014) The bitter taste receptor t2r38 is an independent risk factor for chronic rhinosinusitis requiring sinus surgery. Int Forum Allergy Rhinol 4:3-7 doi: 10.1002/alr.21253

14 Adappa ND, Farquhar, D, Palmer, JN, Kennedy, DW, Doghramji, L, Morris, SA, Owens, D, Mansfield, C, Lysenko, A, Lee, RJ, Cowart, BJ, Reed, DR and Cohen, NA (2015) Tas2r38 genotype predicts surgical outcome in nonpolypoid chronic rhinosinusitis. Int Forum Allergy Rhinol 6:25-33 doi: 10.1002/alr.21666

15 Mfuna Endam L, Filali-Mouhim, A, Boisvert, P, Boulet, LP, Bosse, Y and Desrosiers, M (2014) Genetic variations in taste receptors are associated with chronic rhinosinusitis: A replication study. Int Forum Allergy Rhinol 4:200-206 doi: 10.1002/alr.21275

16 Rom DI, Christensen, JM, Alvarado, R, Sacks, R and Harvey, RJ (2017) The impact of bitter taste receptor genetics on culturable bacteria in chronic rhinosinusitis. Rhinology 55:90-94 doi: 10.4193/Rhin16.181

17 Dzaman K, Zagor, M, Sarnowska, E, Krzeski, A and Kantor, I (2016) The correlation of tas2r38 gene variants with higher risk for chronic rhinosinusitis in polish patients. Otolaryngol. Pol. 70:13-18 doi: 10.5604/00306657.1209438

18 Maurer S, Wabnitz, GH, Kahle, NA, Stegmaier, S, Prior, B, Giese, T, Gaida, MM, Samstag, Y and Hansch, GM (2015) Tasting pseudomonas aeruginosa biofilms: Human neutrophils express the bitter receptor t2r38 as sensor for the quorum sensing molecule n-(3-oxododecanoyl)-l-homoserine lactone. Front. Immunol. 6:369 doi: 10.3389/fimmu.2015.00369

19 Gaida MM, Dapunt, U and Hansch, GM (2016) Sensing developing biofilms: The bitter receptor t2r38 on myeloid cells. Pathog Dis 74:pii: ftw004 doi: 10.1093/femspd/ftw004

20 Tran HTT, Herz, C, Ruf, P, Stetter, R and Lamy, E (2018) Human t2r38 bitter taste receptor expression in resting and activated lymphocytes. Front. Immunol. 9:2949 doi: 10.3389/fimmu.2018.02949

21 Malki A, Fiedler, J, Fricke, K, Ballweg, I, Pfaffl, MW and Krautwurst, D (2015) Class i odorant receptors, tas1r and tas2r taste receptors, are markers for subpopulations of circulating leukocytes. J. Leukoc. Biol. 97:533-545 doi: 10.1189/jlb.2A0714-331RR

22 Bogdan C (2015) Nitric oxide synthase in innate and adaptive immunity: An update. Trends Immunol. 36:161-178 doi: 10.1016/j.it.2015.01.003

23 Huang Z, Hoffmann, FW, Fay, JD, Hashimoto, AC, Chapagain, ML, Kaufusi, PH and Hoffmann, PR (2012) Stimulation of unprimed macrophages with immune complexes triggers a low output of nitric oxide by calcium-dependent neuronal nitric-oxide synthase. J. Biol. Chem. 287:4492-4502 doi: 10.1074/jbc.M111.315598

24 Reiling N, Ulmer, AJ, Duchrow, M, Ernst, M, Flad, HD and Hauschildt, S (1994) Nitric oxide synthase: Mrna expression of different isoforms in human monocytes/macrophages. Eur. J. Immunol. 24:1941-1944 doi: 10.1002/eji.1830240836

25 Mühl H and Pfeilschifter, J (2003) Endothelial nitric oxide synthase: A determinant of tnfα production by human monocytes/macrophages. Biochem. Biophys. Res. Commun. 310:677-680 doi: 10.1016/j.bbrc.2003.09.039

26 Dugas B, Paul-Eugene, N, Yamaoka, K, Amirand, C, Damais, C and Kolb, JP (1995) Il-4 induces camp and cgmp in human monocytic cells. Mediators Inflamm. 4:298-305 doi: 10.1155/S0962935195000482

27 Schmidt HH, Warner, TD, Nakane, M, Forstermann, U and Murad, F (1992) Regulation and subcellular location of nitrogen oxide synthases in raw264.7 macrophages. Mol. Pharmacol. 41:615-624

28 Connelly L, Jacobs, AT, Palacios-Callender, M, Moncada, S and Hobbs, AJ (2003) Macrophage endothelial nitric-oxide synthase autoregulates cellular activation and pro-inflammatory protein expression. J. Biol. Chem. 278:26480-26487 doi: 10.1074/jbc.M302238200

29 Fernandez-Boyanapalli R, McPhillips, KA, Frasch, SC, Janssen, WJ, Dinauer, MC, Riches, DW, Henson, PM, Byrne, A and Bratton, DL (2010) Impaired phagocytosis of apoptotic cells by macrophages in chronic granulomatous disease is reversed by ifn-gamma in a nitric oxide-dependent manner. J. Immunol. 185:4030-4041 doi: 10.4049/jimmunol.1001778

30 Freemerman AJ, Johnson, AR, Sacks, GN, Milner, JJ, Kirk, EL, Troester, MA, Macintyre, AN, Goraksha-Hicks, P, Rathmell, JC and Makowski, L (2014) Metabolic reprogramming of macrophages: Glucose transporter 1 (glut1)-mediated glucose metabolism drives a proinflammatory phenotype. J. Biol. Chem. 289:7884-7896 doi: 10.1074/jbc.M113.522037

31 Wiener A, Shudler, M, Levit, A and Niv, MY (2012) Bitterdb: A database of bitter compounds. Nucleic Acids Res. 40:D413-419 doi: 10.1093/nar/gkr755

32 Meyerhof W, Batram, C, Kuhn, C, Brockhoff, A, Chudoba, E, Bufe, B, Appendino, G and Behrens, M (2010) The molecular receptive ranges of human tas2r bitter taste receptors. Chem. Senses 35:157-170 doi: bjp092 [pii] 10.1093/chemse/bjp092

33 Zhang CH, Lifshitz, LM, Uy, KF, Ikebe, M, Fogarty, KE and ZhuGe, R (2013) The cellular and molecular basis of bitter tastant-induced bronchodilation. PLoS Biol. 11:e1001501 doi: 10.1371/journal.pbio.1001501

34 Roland WS, Gouka, RJ, Gruppen, H, Driesse, M, van Buren, L, Smit, G and Vincken, JP (2014) 6-methoxyflavanones as bitter taste receptor blockers for htas2r39. PLoS One 9:e94451 doi: 10.1371/journal.pone.0094451

35 Greene TA, Alarcon, S, Thomas, A, Berdougo, E, Doranz, BJ, Breslin, PA and Rucker, JB (2011) Probenecid inhibits the human bitter taste receptor tas2r16 and suppresses bitter perception of salicin. PLoS One 6:e20123 doi: 10.1371/journal.pone.0020123

36 Tewson PH, Martinka, S, Shaner, NC, Hughes, TE and Quinn, AM (2016) New dag and camp sensors optimized for live-cell assays in automated laboratories. J Biomol Screen 21:298-305 doi: 10.1177/1087057115618608

37 Klarenbeek J, Goedhart, J, van Batenburg, A, Groenewald, D and Jalink, K (2015) Fourth-generation epac-based fret sensors for camp feature exceptional brightness, photostability and dynamic range: Characterization of dedicated sensors for flim, for ratiometry and with high affinity. PLoS One 10:e0122513 doi: 10.1371/journal.pone.0122513

38 Depry C, Allen, MD and Zhang, J (2011) Visualization of pka activity in plasma membrane microdomains. Mol. Biosyst. 7:52-58 doi: 10.1039/c0mb00079e

39 O’Neill LA and Pearce, EJ (2016) Immunometabolism governs dendritic cell and macrophage function. J. Exp. Med. 213:15-23 doi: 10.1084/jem.20151570

40 Mayevsky A and Rogatsky, GG (2007) Mitochondrial function in vivo evaluated by nadh fluorescence: From animal models to human studies. Am. J. Physiol. Cell Physiol. 292:C615-640 doi: 10.1152/ajpcell.00249.2006

41 Rossi AG, McCutcheon, JC, Roy, N, Chilvers, ER, Haslett, C and Dransfield, I (1998) Regulation of macrophage phagocytosis of apoptotic cells by camp. J. Immunol. 160:3562-3568

42 Yeager LA, Chopra, AK and Peterson, JW (2009) Bacillus anthracis edema toxin suppresses human macrophage phagocytosis and cytoskeletal remodeling via the protein kinase a and exchange protein activated by cyclic amp pathways. Infect. Immun. 77:2530-2543 doi: 10.1128/IAI.00905-08

43 Peters-Golden M (2009) Putting on the brakes: Cyclic amp as a multipronged controller of macrophage function. Sci Signal 2:pe37 doi: 10.1126/scisignal.275pe37

44 Aronoff DM, Canetti, C, Serezani, CH, Luo, M and Peters-Golden, M (2005) Cutting edge: Macrophage inhibition by cyclic amp (camp): Differential roles of protein kinase a and exchange protein directly activated by camp-1. J. Immunol. 174:595-599 doi: 10.4049/jimmunol.174.2.595

45 Townsend EA, Meuchel, LW, Thompson, MA, Pabelick, CM and Prakash, YS (2011) Estrogen increases nitric-oxide production in human bronchial epithelium. J. Pharmacol. Exp. Ther. 339:815-824 doi: 10.1124/jpet.111.184416

46 Lee RJ, Chen, B, Redding, KM, Margolskee, RF and Cohen, NA (2014) Mouse nasal epithelial innate immune responses to pseudomonas aeruginosa quorum-sensing molecules require taste signaling components. Innate Immun. 20:606-617 doi: 10.1177/1753425913503386

47 Roland WS, van Buren, L, Gruppen, H, Driesse, M, Gouka, RJ, Smit, G and Vincken, JP (2013) Bitter taste receptor activation by flavonoids and isoflavonoids: Modeled structural requirements for activation of htas2r14 and htas2r39. J. Agric. Food Chem. 61:10454-10466 doi: 10.1021/jf403387p

48 Kuroda Y, Ikeda, R, Yamazaki, T, Ito, K, Uda, K, Wakabayashi, K and Watanabe, T (2016) Activation of human bitter taste receptors by polymethoxylated flavonoids. Biosci. Biotechnol. Biochem. 80:2014-2017 doi: 10.1080/09168451.2016.1184558

49 Behrens M, Gu, M, Fan, S, Huang, C and Meyerhof, W (2017) Bitter substances from plants used in traditional chinese medicine exert biased activation of human bitter taste receptors. Chem. Biol. Drug Des. 91:422-433 doi: 10.1111/cbdd.13089

50 Levit A, Nowak, S, Peters, M, Wiener, A, Meyerhof, W, Behrens, M and Niv, MY (2014) The bitter pill: Clinical drugs that activate the human bitter taste receptor tas2r14. FASEB J. 28:1181-1197 doi: 10.1096/fj.13-242594

51 Clark AA, Liggett, SB and Munger, SD (2012) Extraoral bitter taste receptors as mediators of off-target drug effects. FASEB J. 26:4827-4831 doi: 10.1096/fj.12-215087

52 Jun CD, Han, MK, Kim, UH and Chung, HT (1996) Nitric oxide induces adp-ribosylation of actin in murine macrophages: Association with the inhibition of pseudopodia formation, phagocytic activity, and adherence on a laminin substratum. Cell. Immunol. 174:25-34 doi: 10.1006/cimm.1996.0290

53 Ke X, Terashima, M, Nariai, Y, Nakashima, Y, Nabika, T and Tanigawa, Y (2001) Nitric oxide regulates actin reorganization through cgmp and ca(2+)/calmodulin in raw 264.7 cells. Biochim. Biophys. Acta 1539:101-113 doi: 10.1016/s0167-4889(01)00090-8

54 Liao WT, You, HL, Li, C, Chang, JG, Chang, SJ and Chen, CJ (2015) Cyclic gmp-dependent protein kinase ii is necessary for macrophage m1 polarization and phagocytosis via toll-like receptor 2. J. Mol. Med. (Berl.) 93:523-533 doi: 10.1007/s00109-014-1236-0

55 Heinloth A, Brune, B, Fischer, B and Galle, J (2002) Nitric oxide prevents oxidised ldl-induced p53 accumulation, cytochrome c translocation, and apoptosis in macrophages via guanylate cyclase stimulation. Atherosclerosis 162:93-101

56 Kiemer AK, Hartung, T and Vollmar, AM (2000) Cgmp-mediated inhibition of tnf-alpha production by the atrial natriuretic peptide in murine macrophages. J. Immunol. 165:175-181

57 Handa P, Tateya, S, Rizzo, NO, Cheng, AM, Morgan-Stevenson, V, Han, CY, Clowes, AW, Daum, G, O’Brien, KD, Schwartz, MW, Chait, A and Kim, F (2011) Reduced vascular nitric oxide-cgmp signaling contributes to adipose tissue inflammation during high-fat feeding. Arterioscler. Thromb. Vasc. Biol. 31:2827-2835 doi: 10.1161/ATVBAHA.111.236554

58 Tateya S, Rizzo, NO, Handa, P, Cheng, AM, Morgan-Stevenson, V, Daum, G, Clowes, AW, Morton, GJ, Schwartz, MW and Kim, F (2011) Endothelial no/cgmp/vasp signaling attenuates kupffer cell activation and hepatic insulin resistance induced by high-fat feeding. Diabetes 60:2792-2801 doi: 10.2337/db11-0255

59 Metukuri MR, Namas, R, Gladstone, C, Clermont, T, Jefferson, B, Barclay, D, Hermus, L, Billiar, TR, Zamora, R and Vodovotz, Y (2009) Activation of latent transforming growth factor-beta1 by nitric oxide in macrophages: Role of soluble guanylate cyclase and map kinases. Wound Repair Regen. 17:578-588 doi: 10.1111/j.1524-475X.2009.00509.x

60 Von Knethen A and Brune, B (2002) Activation of peroxisome proliferator-activated receptor gamma by nitric oxide in monocytes/macrophages down-regulates p47phox and attenuates the respiratory burst. J. Immunol. 169:2619-2626

61 Brune B, Dehne, N, Grossmann, N, Jung, M, Namgaladze, D, Schmid, T, von Knethen, A and Weigert, A (2013) Redox control of inflammation in macrophages. Antioxid Redox Signal 19:595-637 doi: 10.1089/ars.2012.4785

62 Takami S, Getchell, TV, McLaughlin, SK, Margolskee, RF and Getchell, ML (1994) Human taste cells express the g protein alpha-gustducin and neuron-specific enolase. Brain Res. Mol. Brain Res. 22:193-203

63 McLaughlin SK, McKinnon, PJ and Margolskee, RF (1992) Gustducin is a taste-cell-specific g protein closely related to the transducins. Nature 357:563-569 doi: 10.1038/357563a0

64 Foskett JK, White, C, Cheung, KH and Mak, DO (2007) Inositol trisphosphate receptor ca2+ release channels. Physiol. Rev. 87:593-658

65 Yule DI, Straub, SV and Bruce, JI (2003) Modulation of ca2+ oscillations by phosphorylation of ins(1,4,5)p3 receptors. Biochem. Soc. Trans. 31:954-957

66 Ma Z, Taruno, A, Ohmoto, M, Jyotaki, M, Lim, JC, Miyazaki, H, Niisato, N, Marunaka, Y, Lee, RJ, Hoff, H, Payne, R, Demuro, A, Parker, I, Mitchell, CH, Henao-Mejia, J, Tanis, JE, Matsumoto, I, Tordoff, MG and Foskett, JK (2018) Calhm3 is essential for rapid ion channel-mediated purinergic neurotransmission of gpcr-mediated tastes. Neuron 98:547-561 e510 doi: 10.1016/j.neuron.2018.03.043

67 McMahon DB, Workman, AD, Kohanski, MA, Carey, RM, Freund, JR, Hariri, BM, Chen, B, Doghramji, LJ, Adappa, ND, Palmer, JN, Kennedy, DW and Lee, RJ (2018) Protease-activated receptor 2 activates airway apical membrane chloride permeability and increases ciliary beating. FASEB J. 32:155-167 doi: 10.1096/fj.201700114RRR

68 Lee RJ, Hariri, BM, McMahon, DB, Chen, B, Doghramji, L, Adappa, ND, Palmer, JN, Kennedy, DW, Jiang, P, Margolskee, RF and Cohen, NA (2017) Bacterial d-amino acids suppress sinonasal innate immunity through sweet taste receptors in solitary chemosensory cells. Sci Signal 10 doi: 10.1126/scisignal.aam7703

