## Supplementary Material for "Bitter taste receptors stimulate phagocytosis in human macrophages through calcium, nitric oxide, and cyclic-GMP signaling"

### Supplementary Methods

*Reagents and solutions*

Fura-2-acetoxymethyl ester (AM), fluo-4-AM, oregon green 488 BAPTA-1-AM, and DAF-FM diacetate, T2R4 (OSR00153W) and T2R46 (OSR00137W) antibodies were from Thermo Fisher Scientific (Waltham, MA USA). T2R16 (ab75106) and GLUT1 (ab15039) antibodies were from Abcam. BAPTA-AM, parthenolide, apigenin, L- and D-$N^G$-nitroarginine methyl ester (L-NAME and D-NAME), cPTIO, xestospongin C, thapsigargin, ionomycin, U73122, U73343, KT5823, ranitidine, cetirizine, 3oxoC12HSL, and fluorometric $NO_2^-$/$NO_3^-$ kit were from Cayman (Ann Arbor, MI USA). 4'-fluoro-6-methoxyflavanone (2-(4-fluorophenyl)-6-methoxychroman-4-one) was purchased from VitaScreen, LLC (Urbana-Champaign, IL USA). Plasmid encoding AKAR-4 [1] (Addgene # 61619) was from Jin Zhang (University of California San Diego, San Diego, CA USA). EPAC-$S^{H187}$ construct (mTurq2Δ_Epac(CD,ΔDEP, Q270E)_td$^{cp173}$Ven) [2] was from K. Jalink and J. Klarenbeek (Netherlands Cancer Institute, Amsterdam, The Netherlands). Recombinant human IL-4 and M-CSF and ELISAs for IL-10 and IL-12 were from Peprotech (Rocky Hill, NJ USA). Pertussis toxin (PTX) was from Tocris (Minneapolis, MN USA) Unless indicated elsewhere, all other reagents were from Sigma Aldrich (St. Louis, MO USA).

Stock solutions of PQS, HHQ, DHQ, and 3oxoC12HSL were made at 100 mM in DMSO (≥1000x). PQS, HHQ, and 3oxoC12HSL have poor aqueous solubility that may be enhanced *in vivo* biosurfactant rhamnolipids produced by these bacteria [3]. We noted a precipitation and loss of activity of PQS and HHQ after ~30 min in aqueous solution; therefore, working solutions were diluted from stocks immediately before use for each experiment with vigorous vortexing (>90 sec) prior to addition of a 2x working solution (200 µM PQS or HHQ for most experiments) of the compound into a well of a chambered coverglass (CellVis, Mountain View, CA USA) containing cells and an equal volume of HBSS.

*Immunofluorescence (IF) microscopy*

Macrophages (MΦs) were fixed in 4% formaldehyde in DPBS for 20 min at room temperature, followed by blocking and permeabilization in phosphate-buffered saline (PBS) containing 1% bovine serum albumin (BSA), 5% normal donkey serum (NDS), 0.2% saponin, and 0.1% triton X-100 for 1 hour at 4°C. Primary antibody incubation (1:100 for anti-T2R antibodies, 1:250 for Glut1 antibody) were carried out at 4°C overnight. AlexaFluor-labeled donkey anti-mouse or rabbit secondary antibody incubation (1:1000) was carried out for 2 hours at 4°C. Cells were washed and mounted with Fluoroshield with DAPI (Abcam). Images were taken on an Olympus DSU spinning disk confocal system with IX-83 microscope (Olympus Life Sciences, Tokyo, Japan) and 60x (1.4 NA) objective with Metamorph (Molecular Devices, Sunnyvale, CA USA). Images were analyzed using FIJI [4].

*MΦ siRNA reagents and protocols*

Accell SMARTpool siRNAs (Dharmacon, Lafayette, CO USA) designed for primary cells were used at 1 μM final concentration. Stock solutions of siRNA pools were made at 100 μM in siRNA buffer (Cat # B-002000-UB-100) and final working delivery solution was made in Accell Delivery media (Cat # B-005000) containing 2.5% human serum and 1x pen/strep. Media was changed to siRNA media at day 7 and fresh siRNA media was added at day 10, followed by use at day 12 (total ~120 hours exposure to siRNA). Some experiments used ON-TARGET plus SMARTpool siRNAs (Dharmacon, Lafayette, CO USA), which were transfected with DharmaFECT transfection reagent at 0.2 μM final concentration. Media was changed to serum-free RPMI 1640 + siRNAs + DharmaFECT for 6 hours at day 10, followed by replacement with serum-containing media for ~18 hrs. On day 11, another 6 hour pulse of siRNAs in serum-free media was delivered, followed by use at day 13.

*Live cell imaging of intracellular calcium and reactive nitrogen species (RNS) production*

MΦs were loaded with 5 μM of acetoxymethyl ester (AM) variant of the indicator dye for 45 min at room temperature/air in HEPES-buffered HBSS followed by washing with HBSS to remove unloaded dye and 20 min incubation in the dark. Imaging of fura-2 was performed using an Olympus IX-83 microscope (20x 0.75 NA PlanApo objective) equipped with a fluorescence xenon lamp (Sutter Lambda LS, Sutter Instruments, Novato, CA USA), excitation and emission filter wheels (Sutter Lambda 2), and Orca Flash 4.0 sCMOS camera (Hamamatsu, Tokyo, Japan). Images were acquired with MetaFluor (Molecular Devices, Sunnyvale, CA USA) using a standard fura-2 dual excitation filter set (79002-ET, Chroma Technologies, Rockingham, VT USA). Excitation of DAF-FM was carried out with 470/40 nm excitation filter, 495 lp dichroic, and 525/40 nm em filter (49002-ET, Chroma Technologies).

Fluo-4 experiments were performed on a Nikon TS100 microscope with a 20x 0.75 PlanApo objective (Nikon Instruments, Tokyo, Japan) using a standard GFP filter set with excitation from an XCite 110 LED (Excelitas Technologies, Waltham MA USA) and emission captured on a Retiga R1 Camera (Teledyne QImaging, Surrey, BC, Canada). Experiments were done in HBSS (+20 mM HEPES) containing 1 x MEM amino acids (Gibco, Gaithersburg MD USA) to provide a physiological source of extracellular L-arginine.

H441 cells were loaded with DAF-FM or calcium indicator Oregon Green BAPTA 488 by incubation with 5 μM DAF-FM diacetate or Oregon Green BAPTA 488-AM for 60 min. Imaging was performed as for DAF-FM or Fluo-4, respectively, as described above. Experiments were performed in HBSS plus 20 mM HEPES containing 1x MEM amino acids (Gibco, Gaithersburg, MD USA) to provide a physiological level of extracellular L-arginine for NO production.

*Live cell imaging of cAMP and cGMP signaling*

MΦs were infected with BacMams containing either green downward cADDis or green downward GENie (Montana Molecular) to image cAMP or cGMP, respectively. MΦs in 8-well chambered coverglasses were given 100 uL fresh media containing 10% human serum. To each well was added 100 µL containing 20 µL BacMam and 2 mM Sodium butyrate in media. After 6 hours, BacMam solution was removed and 200 µL media + 10% serum was added + 2 mM sodium butyrate. Cells were imaged \ 24-48 hrs after infection as described for Fluo-4 above. For AKAR4 or Epac-$S^{H187}$, 8-well chambered coverglass were transfected with 0.5 µg/well plasmid using Effectene (MilliporeSigma, Burlington, MA USA) as described by the manufacture's protocol using serum-containing media. Cells were imaged 48 hrs. after transfection. Transfection efficiencies of plasmids were low (10-20%), but sufficient for single cell imaging experiments. Experiments with both indicators were done in HBSS +20 mM HEPES containing 1 x MEM amino acids.

*Phagocytosis assays imaging*

Fluorescence microscopy quantification of phagocytosed FITC-labeled *E. coli* was carried out with a 60x 1.4 NA objective, Olympus IX83 microscope, standard FITC filter set, Orca Flash 4.0 camera, and Metamorph. Fluorescence was normalized to number of cells counted via DAPI stained nuclei. We found that the incubation times (1-2 hours) recommended in the assay kit based on RAW J744A.1 murine macrophage-like cells were too long for primary MΦs. Phagocytosis appeared to occur much more quickly with primary cells, therefore quantification was done more quickly. For pHrodo-red-labeled *S. aureus,* a standard TRITC filter set was used.

#### Supplementary Table 1. Bitter compounds used in this study

| Compound | Human T2Rs activated (EC[1] in μM) | References |
|---|---|---|
| Apigenin | T2R14 (8), T2R39 (1) | [5-7] |
| Denatonium benzoate | T2R4 (300), T2R8 (1000), T2R10 (3), T2R13 (30), T2R39 (100), T2R43 (300), T2R46 (30), T2R47 (0.03) | [7, 8] |
| Flufenamic acid (FFA) | T2R14 (0.1) | [7, 8] |
| Heptylhydroxyquinolone (HHQ) | T2R14 (ND[2]), possibly others | [9] |
| Niflumic acid (NFA) | T2R14 (5) | [7, 8] |
| 3oxo-C12HSL | T2R10 (ND), T2R14 (ND), T2R38 (ND) | [10-12] |
| Parthenolide | T2R1 (100), T2R4 (30), T2R8 (100), T2R10 (30), T2R14 (3), T2R44 (100), T2R46 (1). | [7, 8] |
| Phenylthiocarbamine (PTC) | T2R38 (0.04) | [7, 8] |
| Pseudomonas quinolone signal (PQS) | T2R4 (ND), 16 (ND), 38 (ND), possibly others | [9] |
| Quinine | T2R4 (10), T2R7 (10), T2R10 (10), T2R14 (10), T2R39 (10), T2R40 (10), T2R43 (10), T2R44 (10), T2R46 (10) | [7, 8] |
| Salicin | T2R16 (90) | [7, 8] |
| Sodium Benzoate | T2R14 (3000), T2R16 (300) | [7, 8] |
| (-)α-Thujone | T2R10 (100), T2R14 (3) | [7, 8] |

[1]Effective concentration (EC), minimal concentration of agonist that elicits a detectible response, largely based on *in vitro* heterologous expression assays.
[2]Compounds with "ND" denote EC not determined

#### Supplementary Figures

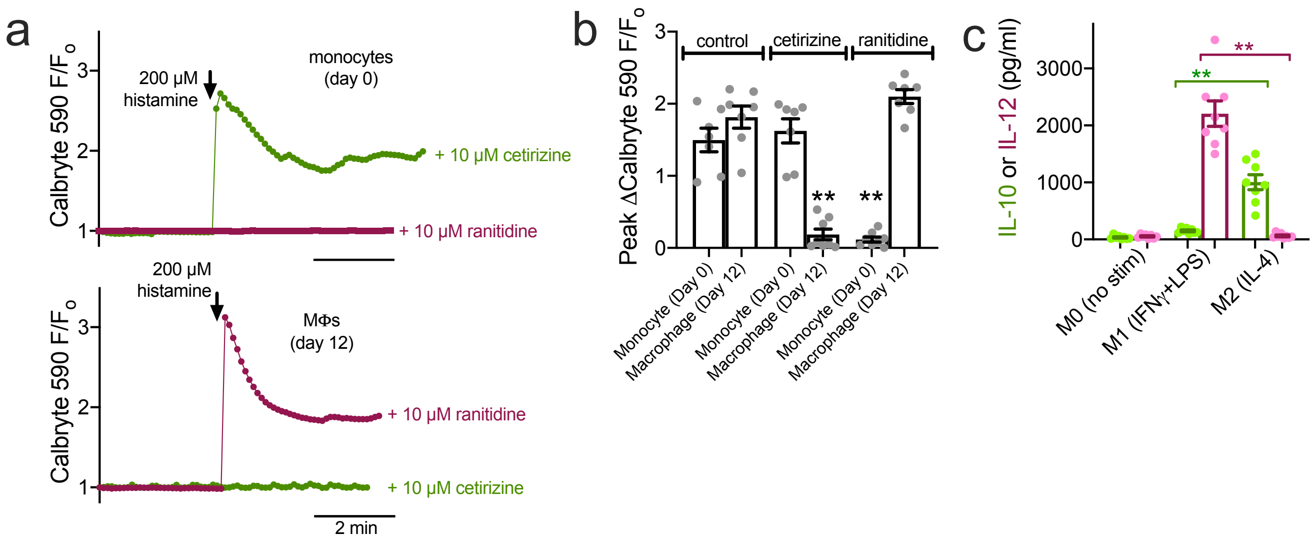


**Supplementary Fig. 1** Confirmation of MΦ differentiation by functional expression of H1 receptors vs H2 receptors, as previously described [13, 14] was determined by calcium imaging. Intracellular calcium was imaged in primary monocytes and MΦs seeded on chambered coverglass. Monocytes were adhered to the glass with CellTak (BD Biosciences) for 30 min. MΦs were differentiated on glass for 10 days in RPMI media containing 10% human serum. Cells washed with HEPES-buffered HBSS and loaded with calcium indicator Calbryte 590-AM (5 µL for 45 min), followed by washing and imaging using a TRITC filter set (Semrock LED-TRITC-A-000) with a 20x 0.75 PlanApo objective (Nikon Instruments, Tokyo Japan) using a standard GFP filter set with excitation from an XCite 110 LED (Excelitas Technologies, Waltham MA USA) and emission captured on a Retiga R1 Camera (Teledyne QImaging, Surrey, BC, Canada). Images were acquired with Micromanager [15]. **a** Representative traces of calcium responses in monocytes (top) and MΦs (after 10 days of adherence differentiation; bottom) in the presence of H1 antagonist cetirizine (green) or H2 antagonist ranitidine (magenta). **b** Bar graph of mean ± SEM of ≥6 independent experiments with histamine stimulation in the absence or presence of cetirizine or ranitidine. Note responses to histamine were inhibited by h1 antagonist cetirizine in MΦs while they were inhibited by H2 antagonist ranitidine in monocytes, supporting the previously described transition from H2 to H1 receptor expression that occurs with differentiation. Significance determined by one-way ANOVA with Bonferonni posttest; ***p*<0.01. **c** As previously described, stimulation of differentiated MΦs with a cytokines designed to promote M1 (20 ng/ml IFNγ + 100 ng/ml LPS [16, 17]) or M2 polarization (20 ng/ml IL-4 [16, 17]) resulted in secretion of M1 marker IL-12 or M2 marker IL-10, as determined by ELISA. Stimulation with LPS/IFNγ or IL-4 was carried out during the final 3 days of differentiation (days 8, 9, and 10). LPS and IFNγ were from Cell Signaling Technologies (Danvers, MA USA) and IL-4 was from Peprotech (Rocky Hill, NJ USA)

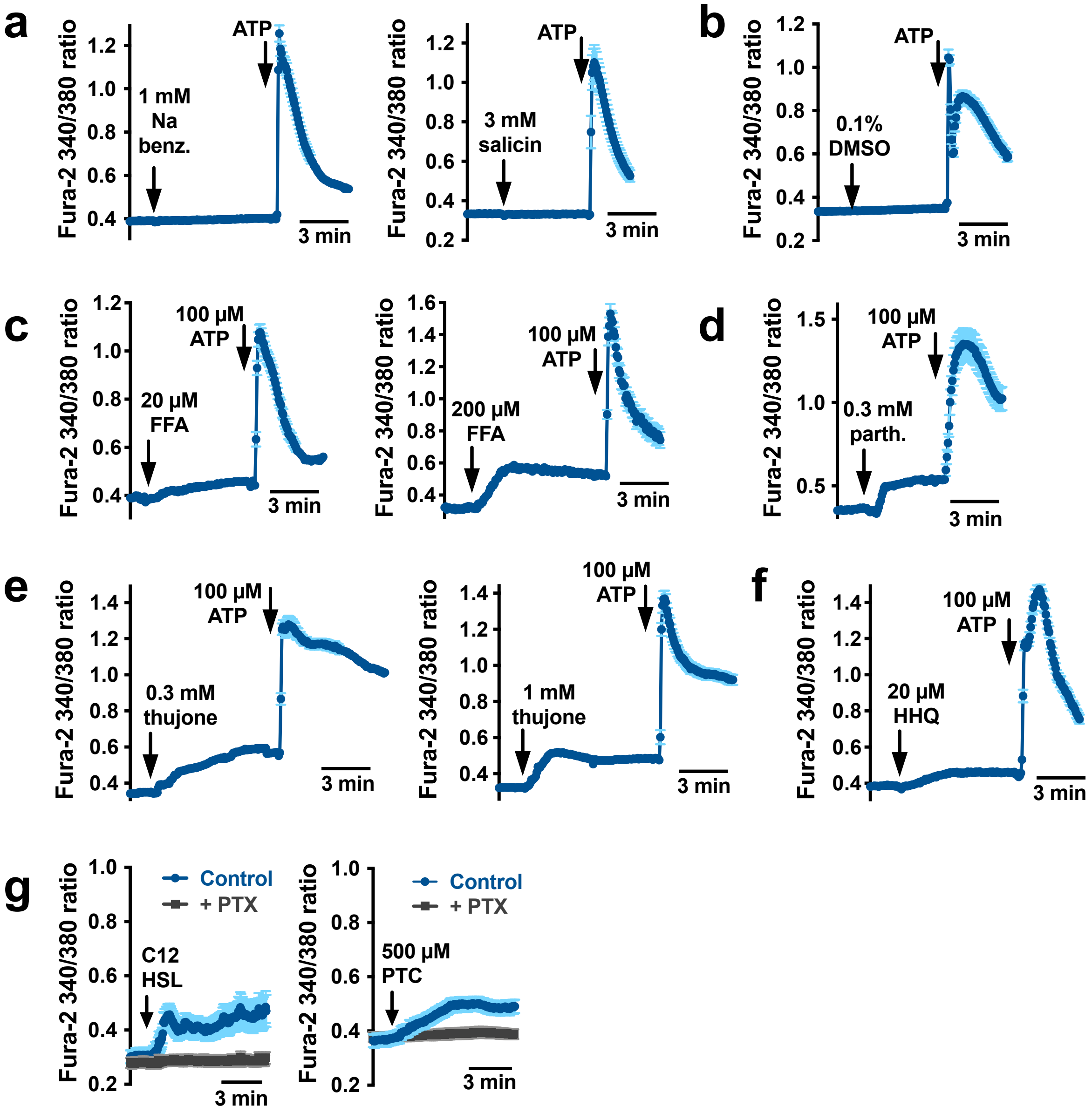


**Supplementary Fig. 2** Representative calcium traces (mean ± SEM) from single experiments of 10-30 MΦs from bar graph data shown in Fig. 1. **a** Representative traces showing no calcium response to NaBenzoate, a weak T2R14 agonist (minimal effective concentration [EC] of 300 µM vs 0.1 µM for FFA in HEK293T heterologous expression assays [7, 8]) and weak T2R16 agonist (EC 3 mM vs 90 µM for salicin [7, 8]). T2R16 agonist salicin also had no effect. 100 µM ATP was used as a positive control to activate purinergic receptors. **b** Vehicle control (0.1% DMSO) had no effect. **c** Representative traces of 20 and 200 µM T2R14 agonist flufenamic acid (FFA). **d** Representative trace of parthenolide, which activates several T2Rs [7, 8]. **e** Trace showing response to T2R10 and T2R14 agonist thujone [7, 8]. **f** Representative trace showing response to T2R14 agonist and *Pseudomonas aeruginosa* quorum-sensing molecule heptylhydroxyquinolone (HHQ [9]). **g** Traces showing responses to T2R agonist 3-oxo-dodecanoyl-homoserine lactone (C12HSL) and T2R38 agonist phenylthiocarbamide (PTC) and inhibition by pertussis toxin (100 ng/ml, 18hrs pretreatment)

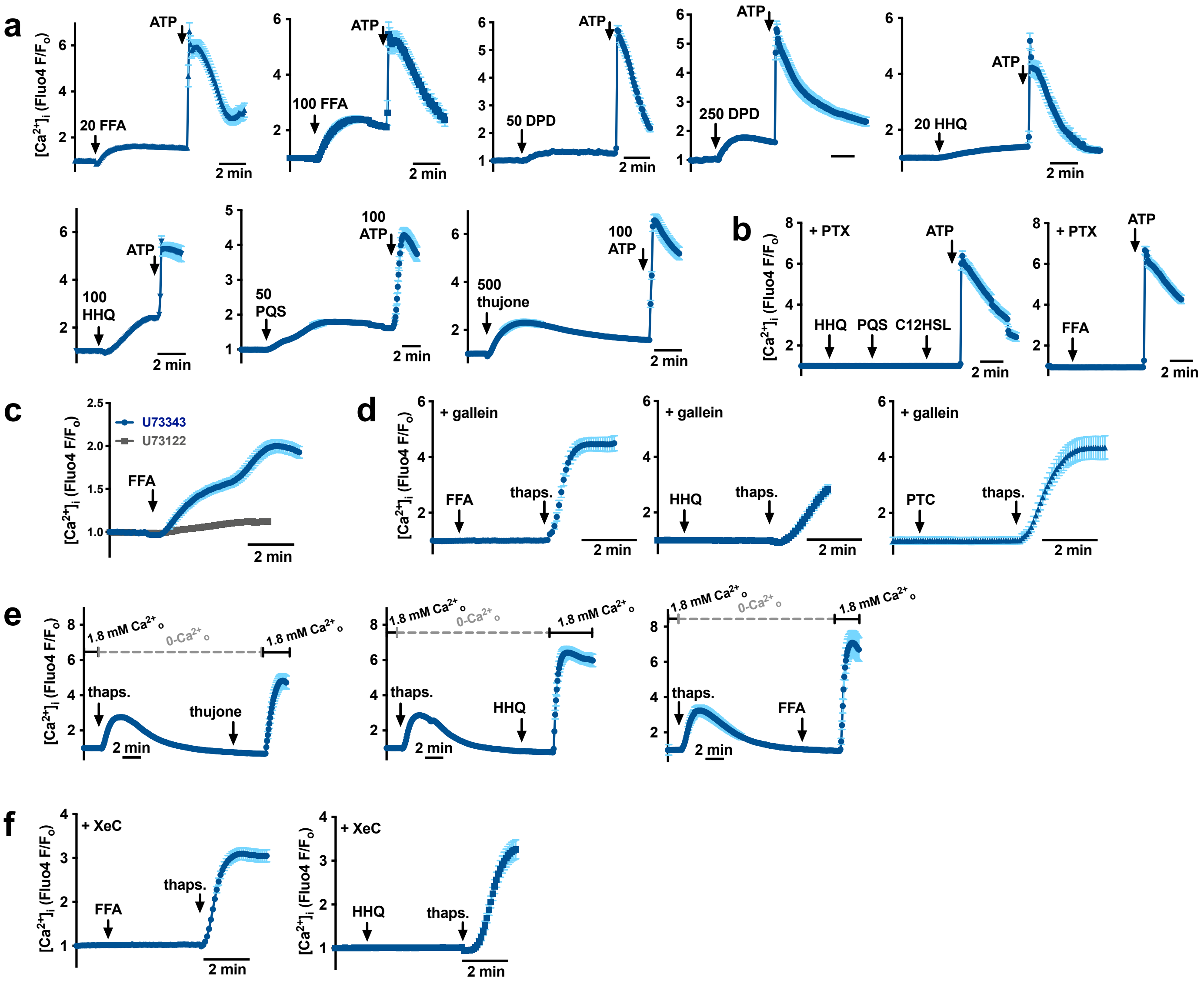

**Supplementary Fig. 3** Representative traces of fluo-4 experiments from Fig. 1. Calcium responses observed using fura-2 were confirmed using fluo-4, and origin of the calcium release examined. **a** Dose dependency of various T2R14 agonists, including flufenamic acid (FFA), diphenhydramine (DPD), heptylhydroxyquinolone (HHQ), *Pseudomonas* quinolone signal (PQS), and thujone. **b** Traces showing inhibition of responses to several compounds from *a* by PTX (100 ng/ml, 18 hrs.). **c** Inhibition of responses to FFA by phospholipase C (PLC) inhibitor U73122 (10 μM, 30 min pretreatment) but not inactive analogue U73343 (10 μM, 30 min pretreatment). **d** Inhibition of responses to 100 μM FFA, HHQ, and PTC by Gβγ inhibitor gallein (100 μM). **e** After ER store depletion with calcium ATPase inhibitor thapsigargin (as previously described [18]), T2R agonists thujone (600 μM), HHQ (100 μM), or FFA (100 μM) had no effect. **f** After treatment with inositol trisphosphate ($IP_3$) receptor ($IP_3R$) inhibitor xestospongin C (XeC; 10 μM, 30 min pretreatment), responses to 100 μM FFA or HHQ were inhibited. Data are summarized in bar graphs in Fig. 1. Together, these data suggest that Gi-coupled T2R activation by bitter compounds results in PLC activation through the Gβγ subunits to stimulate ER calcium release through $IP_3Rs$, as previously reported for airway cells [6, 9, 10, 19] and taste cells [20]

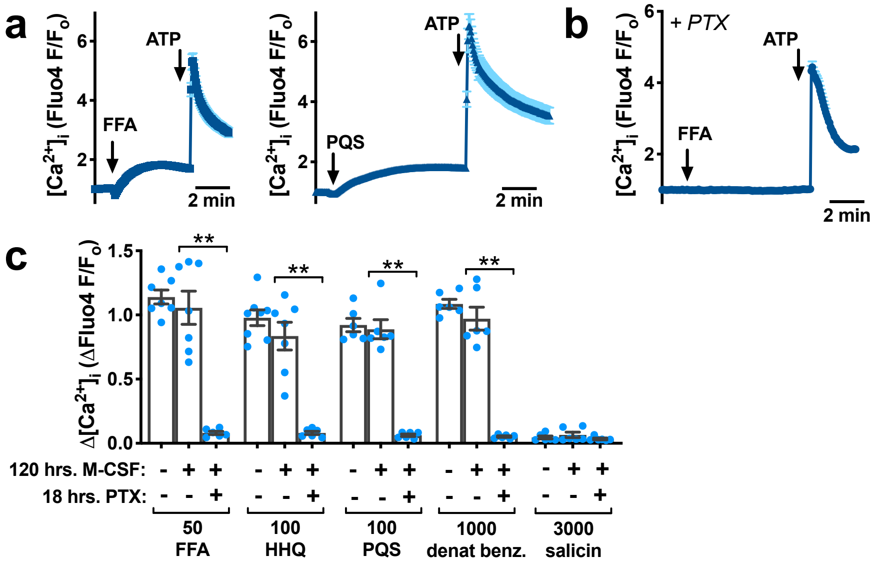


**Supplementary Fig. 4** MΦs differentiated by adherence ± M-CSF (20 ng/ml) showed no significant difference in T2R calcium responses. **a** Representative traces to 50 μM FFA or 100 μM PQS in M-CSF-differentiated MΦs were very similar to MΦs differentiated by adherence only. **b** FFA response were also inhibited by pertussis toxin (PTX; 100 ng/ml 18hrs pretreatment). **c** Responses to T2R agonists (concentrations in μM) showed no significant differences in MΦs ± M-CSF. PTX inhibited responses in M-CSF MΦs as in adherence MΦs (Fig. 1). Significance determined by one way ANOVA with Bonferroni posttest; ***p* <0.01. Other studies have also reported no difference between MΦs differentiated by adherence ± M-CSF for various other parameters [21-23]

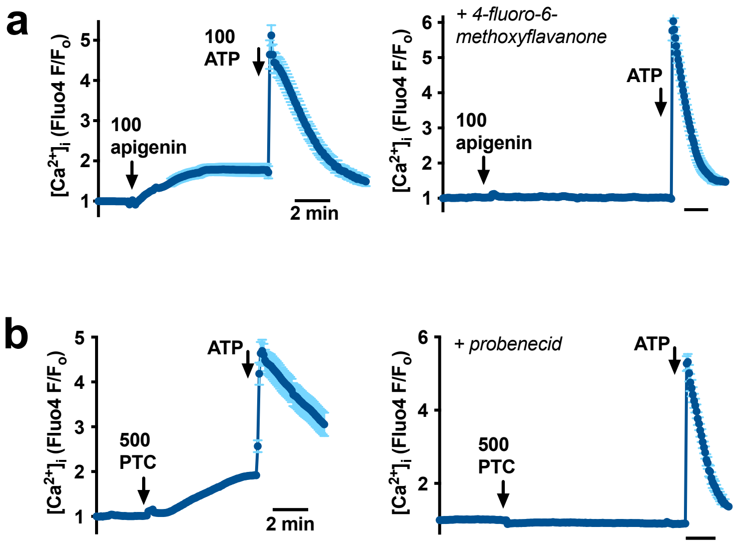


**Supplementary Fig. 5** Inhibition of T2R agonist responses by T2R antagonists. **a** Representative traces from bar graph data in Fig 1 showing inhibition of response to 100 μM apigenin (T2R14/39 agonist) by T2R14/39 antagonist 4-fluoro-6-methoxyflavanone [24] (50 μM, 30 min pretreatment). **b** Representative traces from bar graph data in Fig 1 showing inhibition of response to 500 μM T2R38 agonist PTC by T2R16/38/43 inhibitor probenecid [25]

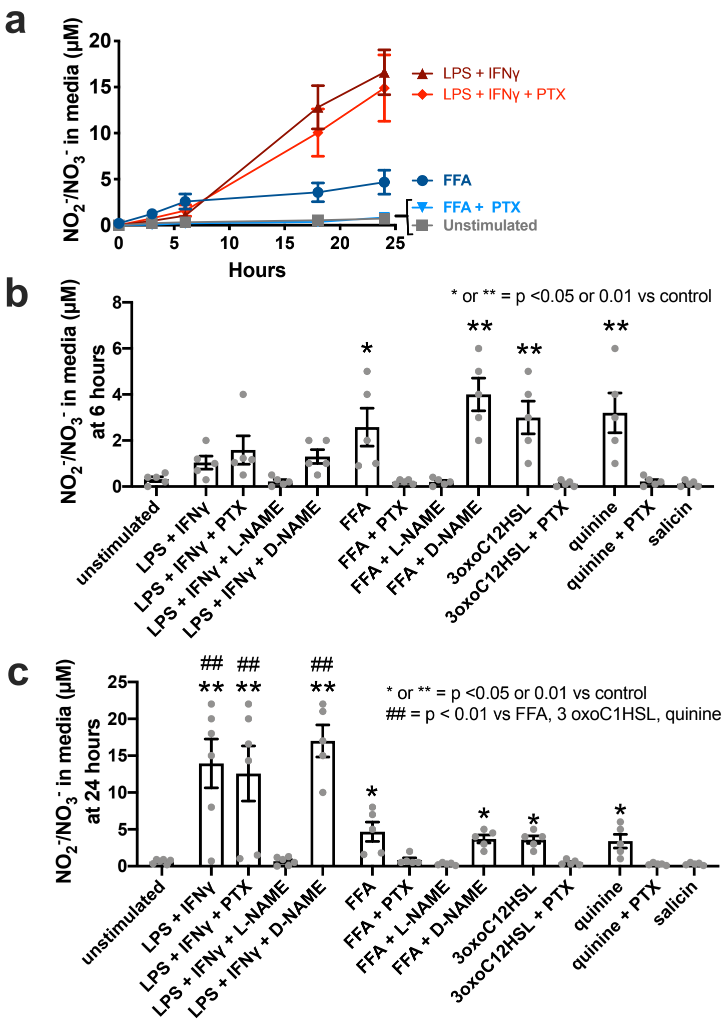


**Supplemental Fig. 6** Confirmation of DAF-FM results with $NO_2^-/NO_3^-$ measurement. As previously done for FcγR-induced e/nNOS activation [26], we measured NO production using a using fluorometric nitric oxide assay kit based on the Greiss reaction (Cayman Chemical) [27]. NO-derived decomposition products ($NO_2^-$ and $NO_3^-$) were measured in media, using untreated media as a control (blank) for baseline $NO_2^-/NO_3^-$. MΦs media was removed at day 12 and replaced with serum-free phenol-red-free DMEM (Gibco) containing stimuli as indicated. Samples of media were collected at various time points as indicated. **a** T2R agonist flufenamic acid (FFA; 100 µM) caused rapid $NO_2^-/NO_3^-$ production rapidly over the first 6 hours that then increased more slowly. In contrast, LPS and IFNγ (as used in Supplementary Fig. 1c) induced a larger $NO_2^-/NO_3^-$ production that occurred more slowly, with a lag, likely reflecting induction of constitutively-active iNOS expression. FFA-induced $NO_2^-/NO_3^-$ production was PTX-sensitive, whereas LPS and IFNγ-induced $NO_2^-/NO_3^-$ production was not. **b-c** Experiments were performed as in *a* and media samples were taken at 6 hrs (*b*) and 24 hrs (*c*). Asterisks represent significance compared with control (unstimulated cultures) by 1-way ANOVA with Dunnett's posttest. Pound signs in *c* indicate significance as indicated determined by 1-way ANOVA with Bonferroni posttest. Note larger $NO_2^-/NO_3^-$ production with T2R agonists at 6 hrs and larger $NO_2^-/NO_3^-$ production with LPS and IFNγ at 24 hrs. T2R-induced $NO_2^-/NO_3^-$ production was PTX-sensitive while LPS and IFNγ-induced $NO_2^-/NO_3^-$ production was not. All $NO_2^-/NO_3^-$ production was inhibited by L-NAME but not D-NAME. Salicin, a T2R16 agonist that did not elicit calcium responses, did not elicit $NO_2^-/NO_3^-$ production. Together, these data support that results from DAF-FM reflect NO production

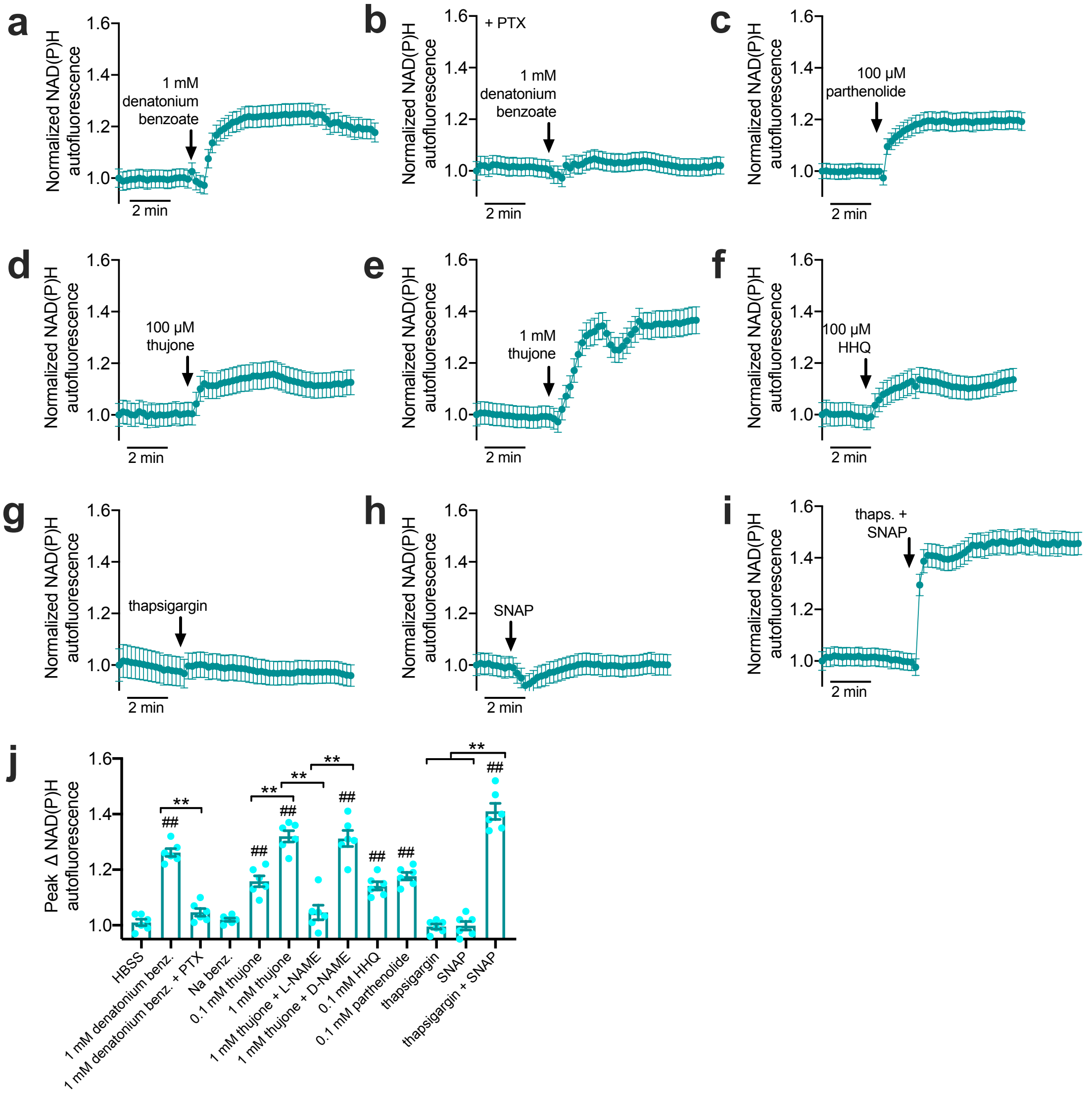


**Supplementary Fig. 7** Imaging changes in NAD(P)H autofluorescence [28, 29] induced by T2R receptor stimulation. This long-established technique utilizes the autofluorescent property of NADH and NADPH, but not NAD and NADP, when excited by UV light [28-30]. We excited MΦs with a Xenon lamp with high UV output at 340 nm and measured fluorescence at 450 nm during stimulation with T2R agonists. Our goal was not to perform a rigorous immuno-metabolic characterization, but rather to test if changes in metabolism could occur during these acute stimulations. MΦs seeded on glass coverslips were washed with HBSS and imaged using a Sutter Lamda LS 300 W lamp, DAPI filter set, and 30x 1.0 NA silicone oil immersion objective lens with high UV transmittance on an IX-83 microscope (Olympus) equipped with a Hammamatsu Orca Flash 4.0 sCMOS camera (Hammamatsu, Tokyo, Japan). Images were acquired every 12 seconds using Metafluor (Molecular Devices, Sunnyvale CA). **a-b** Increase in autofluorescence elicited by 1 mM denatonium benzoate (*a*) and inhibition by pertussis toxin (PTX; 100 ng/ml, 18 hrs pretreatment; *b*). **c-f** Increases in autofluorescence elicited by multi-T2R agonist parthenolide (*c*), T2R10/14 agonist thujone (*d* and *e*) and T2R14 agonist HHQ (*f*). **g-i** Nonspecific elevation of calcium with thapsigargin (10 µg/ml; *g*) or nonspecific NO increase with NO donor S-nitroso-N-acetyl-D,L-penicillamine (SNAP; 10 µM; *h*) did not affect autofluorescence, but the addition of both together did increase autofluorescence (*i*), suggesting this change requires the combined elevation of calcium and NO. **j** Bar graph of peak autofluorescence changes over 5 min with T2R stimulation as indicated. Note inhibition of response to thujone by nitric oxide synthase (NOS) inhibitor L-NAME but not by inactive D-NAME (100 µM; 30 min pretreatment)

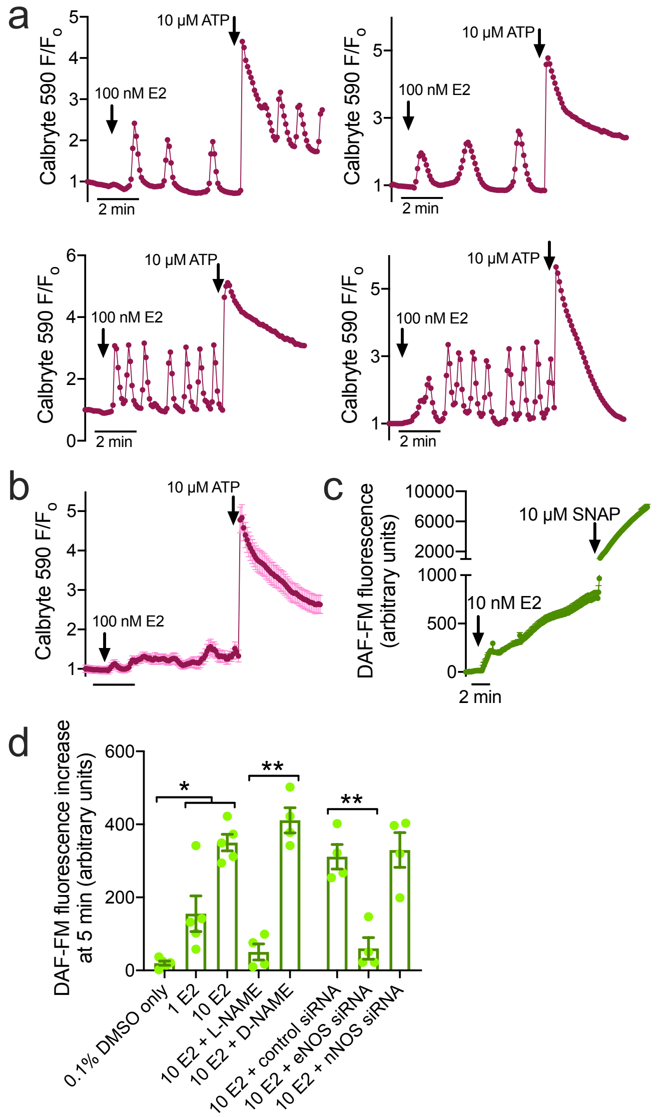


**Supplementary Fig. 8** Calcium signals and NO production induced by 17β-estradiol (E2) in H441 cells. H441 small airway epithelial cells were previously reported to express eNOS [31, 32], as reported for primary airway epithelial cells [33, 34], and H441s produce NO in response to E2 [35], as reported for primary bronchial epithelial cells [36]. Non-genomic acute eNOS activation by E2 is well documented in several tissues [35, 37-43]. H441s were grown on plastic and loaded for 45 min with 5 μM Calbryte 590-AM to visualize calcium responses to 17β-estradiol (E2) using Nikon TS100F with 10x 0.3 NA objective, standard TRITC filter set, Retiga R1 camera (QImaging), and Micromanger [15]. **a** 100 nM E2 resulted in calcium oscillations in individual H441 cells. **b** population average of 30 oscillating cells illustrates an overall sustained low-level calcium elevation. **c** E2 activated NO production as measured using DAF-FM. S-nitroso-N-acetyl-D,L-penicillamine (SNAP) was used as a positive control. Trace represents average of 30 MΦs (mean ± SEM) from one representative experiment. **d** DAF-FM fluorescence increases in response to DMSO (vehicle control), or E2 ± D- or L-NAME; siRNAs (Accell SMARTpool siRNAs; Dharmacon, Lafayette, CO) directed against eNOS or nNOS were also used and suggested that the overwhelming majority of NO production was due eNOS rather than nNOS. Individual points represent individual experiments (n = ≥5, each imaging 20-30 MΦs, MΦs from 2 individuals were used). Bar graph shows mean ± SEM of individual experiments. Significance determined by one-way ANOVA with Bonferroni posttest with pairwise comparisons as indicated; *$p$ <0.05 and **$p$ <0.01

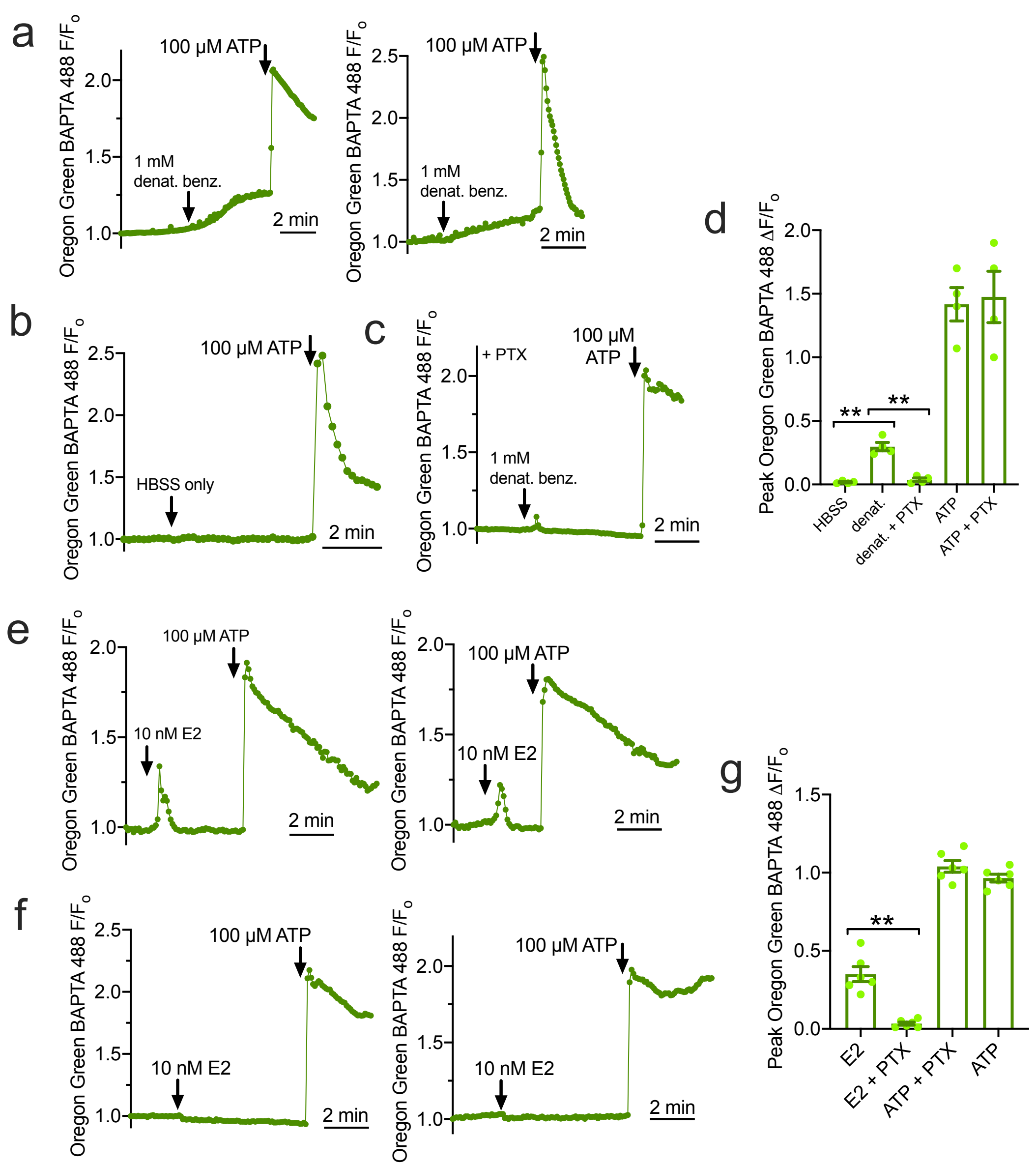


**Supplementary Fig. 9** MΦ calcium responses to 17β-estradiol (E2). **a-d** MΦs loaded with calcium indicator Oregon Green BAPTA 488 (loaded and imaged as described in the text for Fluo-4) exhibited similar low-level responses to denatonium benzoate (*a*) but not HBSS only (vehicle control; *b*) as observed with fluo-4 and fura-2 that were inhibited by pertussis toxin (PTX; 18 hrs 100 ng/ml pretreatment; *c*). ATP responses (driven by $G_q$-coupled purinergic receptors) were intact with PTX (*d*). Oregon Green results validate fluo-4 and fura-2 results in the main text. **e-g** MΦs exhibited low-level calcium transients to 10 nM E2 as previously reported [44, 45] (*e* and *g*), but as also previously reported [44], these were completely blocked by PTX (*f* and *g*). Traces in *a-c* and *e-f* are average of 30 MΦs (mean ± SEM) from one representative experiment for each graph. Bar graphs in *d* and *g* show mean ± SEM from individual experiments (n = ≥5, each imaging 20-30 MΦs, MΦs from 2 individuals were used). Significance determined by one-way ANOVA with Bonferroni posttest with pairwise comparisons as indicated; $^{**}p$ <0.01

#### Supplementary References

1 Depry C, Allen, MD and Zhang, J (2011) Visualization of pka activity in plasma membrane microdomains. Mol. Biosyst. 7:52-58 doi: 10.1039/c0mb00079e

2 Klarenbeek J, Goedhart, J, van Batenburg, A, Groenewald, D and Jalink, K (2015) Fourth-generation epac-based fret sensors for camp feature exceptional brightness, photostability and dynamic range: Characterization of dedicated sensors for flim, for ratiometry and with high affinity. PLoS One 10:e0122513 doi: 10.1371/journal.pone.0122513

3 Calfee MW, Shelton, JG, McCubrey, JA and Pesci, EC (2005) Solubility and bioactivity of the pseudomonas quinolone signal are increased by a pseudomonas aeruginosa-produced surfactant. Infect. Immun. 73:878-882 doi: 10.1128/IAI.73.2.878-882.2005

4 Schindelin J, Arganda-Carreras, I, Frise, E, Kaynig, V, Longair, M, Pietzsch, T, Preibisch, S, Rueden, C, Saalfeld, S, Schmid, B, Tinevez, JY, White, DJ, Hartenstein, V, Eliceiri, K, Tomancak, P and Cardona, A (2012) Fiji: An open-source platform for biological-image analysis. Nat Methods 9:676-682 doi: 10.1038/nmeth.2019

5 Roland WS, van Buren, L, Gruppen, H, Driesse, M, Gouka, RJ, Smit, G and Vincken, JP (2013) Bitter taste receptor activation by flavonoids and isoflavonoids: Modeled structural requirements for activation of htas2r14 and htas2r39. J. Agric. Food Chem. 61:10454-10466 doi: 10.1021/jf403387p

6 Hariri BM, McMahon, DB, Chen, B, Freund, JR, Mansfield, CJ, Doghramji, LJ, Adappa, ND, Palmer, JN, Kennedy, DW, Reed, DR, Jiang, P and Lee, RJ (2017) Flavones modulate respiratory epithelial innate immunity: Anti-inflammatory effects and activation of the t2r14 receptor. J. Biol. Chem. 292:8484-8497 doi: 10.1074/jbc.M116.771949

7 Wiener A, Shudler, M, Levit, A and Niv, MY (2012) Bitterdb: A database of bitter compounds. Nucleic Acids Res. 40:D413-419 doi: 10.1093/nar/gkr755

8 Meyerhof W, Batram, C, Kuhn, C, Brockhoff, A, Chudoba, E, Bufe, B, Appendino, G and Behrens, M (2010) The molecular receptive ranges of human tas2r bitter taste receptors. Chem. Senses 35:157-170 doi: bjp092 [pii] 10.1093/chemse/bjp092

9 Freund JR, Mansfield, CJ, Doghramji, LJ, Adappa, ND, Palmer, JN, Kennedy, DW, Reed, DR, Jiang, P and Lee, RJ (2018) Activation of airway epithelial bitter taste receptors by pseudomonas aeruginosa quinolones modulates calcium, cyclic-amp, and nitric oxide signaling. J. Biol. Chem. 293:9824-9840 doi: 10.1074/jbc.RA117.001005

10 Lee RJ, Xiong, G, Kofonow, JM, Chen, B, Lysenko, A, Jiang, P, Abraham, V, Doghramji, L, Adappa, ND, Palmer, JN, Kennedy, DW, Beauchamp, GK, Doulias, P-T, Ischiropoulos, H, Kreindler, JL, Reed, DR and Cohen, NA (2012) T2r38 taste receptor polymorphisms underlie susceptibility to upper respiratory infection. J. Clin. Invest. 122:4145-4159

11 Lossow K, Hubner, S, Roudnitzky, N, Slack, JP, Pollastro, F, Behrens, M and Meyerhof, W (2016) Comprehensive analysis of mouse bitter taste receptors reveals different molecular receptive ranges for orthologous receptors in mice and humans. J. Biol. Chem. 291:15358-15377 doi: 10.1074/jbc.M116.718544

12 Jaggupilli A, Singh, N, Jesus, VC, Duan, K and Chelikani, P (2018) Characterization of the binding sites for bacterial acyl homoserine lactones (ahls) on human bitter taste receptors (t2rs). ACS Infect Dis 4:1146-1156 doi: 10.1021/acsinfecdis.8b00094

13 Wang KY, Arima, N, Higuchi, S, Shimajiri, S, Tanimoto, A, Murata, Y, Hamada, T and Sasaguri, Y (2000) Switch of histamine receptor expression from h2 to h1 during differentiation of monocytes into macrophages. FEBS Lett. 473:345-348

14 Triggiani M, Petraroli, A, Loffredo, S, Frattini, A, Granata, F, Morabito, P, Staiano, RI, Secondo, A, Annunziato, L and Marone, G (2007) Differentiation of monocytes into macrophages induces the upregulation of histamine h1 receptor. J. Allergy Clin. Immunol. 119:472-481 doi: 10.1016/j.jaci.2006.09.027

15 Edelstein A, Amodaj, N, Hoover, K, Vale, R and Stuurman, N (2010) Computer control of microscopes using micromanager. Curr. Protoc. Mol. Biol. Chapter 14:Unit14 20 doi: 10.1002/0471142727.mb1420s92

16 Vogel DY, Glim, JE, Stavenuiter, AW, Breur, M, Heijnen, P, Amor, S, Dijkstra, CD and Beelen, RH (2014) Human macrophage polarization in vitro: Maturation and activation methods compared. Immunobiology 219:695-703 doi: 10.1016/j.imbio.2014.05.002

17 Murray PJ (2017) Macrophage polarization. Annu. Rev. Physiol. 79:541-566 doi: 10.1146/annurev-physiol-022516-034339

18 McMahon DB, Workman, AD, Kohanski, MA, Carey, RM, Freund, JR, Hariri, BM, Chen, B, Doghramji, LJ, Adappa, ND, Palmer, JN, Kennedy, DW and Lee, RJ (2018) Protease-activated receptor 2 activates airway apical membrane chloride permeability and increases ciliary beating. FASEB J. 32:155-167 doi: 10.1096/fj.201700114RRR

19 Freund JR and Lee, RJ (2018) Taste receptors in the upper airway. World J Otorhinolaryngol Head Neck Surg 4:67-76 doi: 10.1016/j.wjorl.2018.02.004

20 Margolskee RF (2002) Molecular mechanisms of bitter and sweet taste transduction. J. Biol. Chem. 277:1-4 doi: 10.1074/jbc.R100054200

21 Valente RC, Araujo, EG and Rumjanek, VM (2012) Ouabain inhibits monocyte activation in vitro: Prevention of the proinflammatory mcd14(+)/cd16(+) subset appearance and cell-size progression. J. Exp. Pharmacol. 4:125-140 doi: 10.2147/JEP.S35507

22 Lacey DC, Achuthan, A, Fleetwood, AJ, Dinh, H, Roiniotis, J, Scholz, GM, Chang, MW, Beckman, SK, Cook, AD and Hamilton, JA (2012) Defining gm-csf- and macrophage-csf-dependent macrophage responses by in vitro models. J. Immunol. 188:5752-5765 doi: 10.4049/jimmunol.1103426

23 Ohradanova-Repic A, Machacek, C, Fischer, MB and Stockinger, H (2016) Differentiation of human monocytes and derived subsets of macrophages and dendritic cells by the hlda10 monoclonal antibody panel. Clin Transl Immunology 5:e55 doi: 10.1038/cti.2015.39

24 Roland WS, Gouka, RJ, Gruppen, H, Driesse, M, van Buren, L, Smit, G and Vincken, JP (2014) 6-methoxyflavanones as bitter taste receptor blockers for htas2r39. PLoS One 9:e94451 doi: 10.1371/journal.pone.0094451

25 Greene TA, Alarcon, S, Thomas, A, Berdougo, E, Doranz, BJ, Breslin, PA and Rucker, JB (2011) Probenecid inhibits the human bitter taste receptor tas2r16 and suppresses bitter perception of salicin. PLoS One 6:e20123 doi: 10.1371/journal.pone.0020123

26 Huang Z, Hoffmann, FW, Fay, JD, Hashimoto, AC, Chapagain, ML, Kaufusi, PH and Hoffmann, PR (2012) Stimulation of unprimed macrophages with immune complexes triggers a low output of nitric oxide by calcium-dependent neuronal nitric-oxide synthase. J. Biol. Chem. 287:4492-4502 doi: 10.1074/jbc.M111.315598

27 Rassaf T, Feelisch, M and Kelm, M (2004) Circulating no pool: Assessment of nitrite and nitroso species in blood and tissues. Free Radic. Biol. Med. 36:413-422 doi: 10.1016/j.freeradbiomed.2003.11.011

28 Bartolome F and Abramov, AY (2015) Measurement of mitochondrial nadh and fad autofluorescence in live cells. Methods Mol. Biol. 1264:263-270 doi: 10.1007/978-1-4939-2257-4_23

29 Blacker TS and Duchen, MR (2016) Investigating mitochondrial redox state using nadh and nadph autofluorescence. Free Radic. Biol. Med. doi: 10.1016/j.freeradbiomed.2016.08.010

30 Mayevsky A and Rogatsky, GG (2007) Mitochondrial function in vivo evaluated by nadh fluorescence: From animal models to human studies. Am. J. Physiol. Cell Physiol. 292:C615-640 doi: 10.1152/ajpcell.00249.2006

31 Shaul PW, North, AJ, Wu, LC, Wells, LB, Brannon, TS, Lau, KS, Michel, T, Margraf, LR and Star, RA (1994) Endothelial nitric oxide synthase is expressed in cultured human bronchiolar epithelium. J. Clin. Invest. 94:2231-2236 doi: 10.1172/JCI117585

32 German Z, Chambliss, KL, Pace, MC, Arnet, UA, Lowenstein, CJ and Shaul, PW (2000) Molecular basis of cell-specific endothelial nitric-oxide synthase expression in airway epithelium. J. Biol. Chem. 275:8183-8189

33 Sherman TS, Chen, Z, Yuhanna, IS, Lau, KS, Margraf, LR and Shaul, PW (1999) Nitric oxide synthase isoform expression in the developing lung epithelium. Am. J. Physiol. 276:L383-390

34 Stout SL, Wyatt, TA, Adams, JJ and Sisson, JH (2007) Nitric oxide-dependent cilia regulatory enzyme localization in bovine bronchial epithelial cells. J. Histochem. Cytochem. 55:433-442 doi: jhc.6A7089.2007 [pii] 10.1369/jhc.6A7089.2007

35 Kirsch EA, Yuhanna, IS, Chen, Z, German, Z, Sherman, TS and Shaul, PW (1999) Estrogen acutely stimulates endothelial nitric oxide synthase in h441 human airway epithelial cells. Am. J. Respir. Cell Mol. Biol. 20:658-666 doi: 10.1165/ajrcmb.20.4.3241

36 Townsend EA, Meuchel, LW, Thompson, MA, Pabelick, CM and Prakash, YS (2011) Estrogen increases nitric-oxide production in human bronchial epithelium. J. Pharmacol. Exp. Ther. 339:815-824 doi: 10.1124/jpet.111.184416

37 Shaul PW (1999) Rapid activation of endothelial nitric oxide synthase by estrogen. Steroids 64:28-34

38 Wyckoff MH, Chambliss, KL, Mineo, C, Yuhanna, IS, Mendelsohn, ME, Mumby, SM and Shaul, PW (2001) Plasma membrane estrogen receptors are coupled to endothelial nitric-oxide synthase through galpha(i). J. Biol. Chem. 276:27071-27076 doi: 10.1074/jbc.M100312200

39 Wu Q, Chambliss, K, Umetani, M, Mineo, C and Shaul, PW (2011) Non-nuclear estrogen receptor signaling in the endothelium. J. Biol. Chem. 286:14737-14743 doi: 10.1074/jbc.R110.191791

40 Chambliss KL, Yuhanna, IS, Mineo, C, Liu, P, German, Z, Sherman, TS, Mendelsohn, ME, Anderson, RG and Shaul, PW (2000) Estrogen receptor alpha and endothelial nitric oxide synthase are organized into a functional signaling module in caveolae. Circ. Res. 87:E44-52

41 Chambliss KL and Shaul, PW (2002) Estrogen modulation of endothelial nitric oxide synthase. Endocr. Rev. 23:665-686 doi: 10.1210/er.2001-0045

42 Chambliss KL and Shaul, PW (2002) Rapid activation of endothelial no synthase by estrogen: Evidence for a steroid receptor fast-action complex (srfc) in caveolae. Steroids 67:413-419

43 Shaul PW (2002) Regulation of endothelial nitric oxide synthase: Location, location, location. Annu. Rev. Physiol. 64:749-774 doi: 10.1146/annurev.physiol.64.081501.155952

44 Guo Z, Krucken, J, Benten, WP and Wunderlich, F (2002) Estradiol-induced nongenomic calcium signaling regulates genotropic signaling in macrophages. J. Biol. Chem. 277:7044-7050 doi: 10.1074/jbc.M109808200

45 Liu L, Zhao, Y, Xie, K, Sun, X, Gao, Y and Wang, Z (2013) Estrogen-induced nongenomic calcium signaling inhibits lipopolysaccharide-stimulated tumor necrosis factor alpha production in macrophages. PLoS One 8:e83072 doi: 10.1371/journal.pone.0083072
